# Virological characteristics of SARS-CoV-2 BA.2 variant

**DOI:** 10.1101/2022.02.14.480335

**Authors:** Daichi Yamasoba, Izumi Kimura, Hesham Nasser, Yuhei Morioka, Naganori Nao, Jumpei Ito, Keiya Uriu, Masumi Tsuda, Jiri Zahradnik, Kotaro Shirakawa, Rigel Suzuki, Mai Kishimoto, Yusuke Kosugi, Kouji Kobiyama, Teppei Hara, Mako Toyoda, Yuri L Tanaka, Erika P Butlertanaka, Ryo Shimizu, Hayato Ito, Lei Wang, Yoshitaka Oda, Yasuko Orba, Michihito Sasaki, Kayoko Nagata, Kumiko Yoshimatsu, Hiroyuki Asakura, Mami Nagashima, Kenji Sadamasu, Kazuhisa Yoshimura, Jin Kuramochi, Motoaki Seki, Ryoji Fujiki, Atsushi Kaneda, Tadanaga Shimada, Taka-aki Nakada, Seiichiro Sakao, Takuji Suzuki, Takamasa Ueno, Akifumi Takaori-Kondo, Ken J Ishii, Gideon Schreiber, The Genotype to Phenotype Japan (G2P-Japan) Consortium, Hirofumi Sawa, Akatsuki Saito, Takashi Irie, Shinya Tanaka, Keita Matsuno, Takasuke Fukuhara, Terumasa Ikeda, Kei Sato

## Abstract

Soon after the emergence and global spread of a new severe acute respiratory syndrome coronavirus 2 (SARS-CoV-2) Omicron lineage, BA.1 (ref^1, 2^), another Omicron lineage, BA.2, has initiated outcompeting BA.1. Statistical analysis shows that the effective reproduction number of BA.2 is 1.4-fold higher than that of BA.1. Neutralisation experiments show that the vaccine-induced humoral immunity fails to function against BA.2 like BA.1, and notably, the antigenicity of BA.2 is different from BA.1. Cell culture experiments show that BA.2 is more replicative in human nasal epithelial cells and more fusogenic than BA.1. Furthermore, infection experiments using hamsters show that BA.2 is more pathogenic than BA.1. Our multiscale investigations suggest that the risk of BA.2 for global health is potentially higher than that of BA.1.

## Introduction

Virological characteristics of newly emerging SARS-CoV-2 variants, such as transmissibility, pathogenicity and resistance to the vaccine-induced immunity and antiviral drugs, is an urgent global health concern. In February 2022, the Omicron variant (B.1.1.529 and BA lineages) spreads worldwide and represents the most recently recognised variant of concern^2^. Omicron was first reported from South Africa at the end of November 2021^1^. Then, a variant of Omicron, the BA.1 lineage, has rapidly spread worldwide and outcompeted other variants such as Delta.

As of February 2022, another variant of Omicron, the BA.2 lineage, has detected in multiple countries such as Denmark and UK^3^. Notably, BA.2 has initiated outcompeting BA.1^3^, suggesting that BA.2 is more transmissible than BA.1.

In a few months since the emergence of BA.1, we and others revealed the virological characteristics of BA.1^4–19^. For instance, BA.1 is highly resistant to the vaccine-induced humoral immunity and antiviral drugs^4–11, 16–19^. Also, the spike (S) protein of BA.1 is less efficiently cleaved by furin and less fusogenic than those of Delta and an ancestral SARS-CoV-2 belonging to the B.1.1 lineage^11, 12^. Further, the pathogenicity of BA.1 is attenuated when compared to Delta and an ancestral B.1.1 virus^12–14^. However, the virological characteristics of BA.2 remains unaddressed.

### Phylogenetic and epidemic dynamics of BA.2

As of February 2022, Omicron is classified into three main lineages, BA.1, BA.2, BA.3, and a sublineage of BA.1, BA1.1, which harbours the R346K substitution in S (**Fig. 1a**). Although these lineages are monophyletic, their sequences have been greatly diversified. For example, BA.1 differs from BA.2 by 50 amino acids, which is approximately twice as much as the numbers of amino acid differences between four VOCs (Alpha, Beta, Gamma and Delta) and Wuhan-Hu-1, a prototypic SARS-CoV-2 isolate (**Fig. 1b**). Phylodynamics analysis suggests that BA.1 emerged first, followed by BA.2 and BA.3 (**Extended Data Fig. 1**). In addition to BA.1, the earlier strains of BA.2, BA.3, and BA.1.1 were isolated from Gauteng Province, South Africa, the place of the earliest Omicron (BA.1) epidemic (**Extended Data Fig. 1**)^20^. These results suggest that the remarkable diversification of Omicron occurred in Gauteng Province and all Omicron lineages emerged there.

**Fig. 1.**
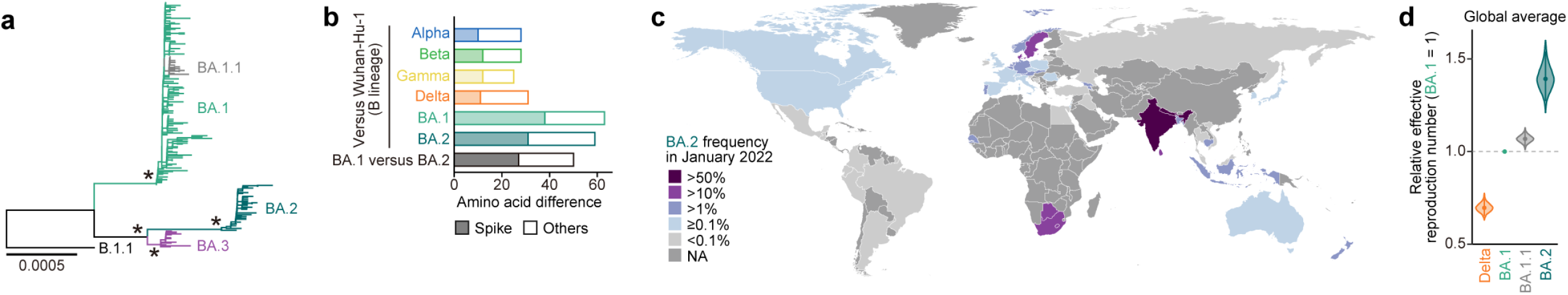
Epidemic of BA.2. **a**, Maximum likelihood tree of the Omicron lineages sampled from South Africa. Asterisk denote nodes with ≥0.95 bootstrap values. **b**, Number of amino acid differences between viral lineages detected in S (filled) or the other regions (opened). **c**, Relative frequency of BA.2 in the genome surveillance data during January 2022. The values for the countries with ≥20 SARS-CoV-2 sequences are shown. **d**, Estimated relative effective reproduction number of each viral lineage, assuming a fixed generation time of 2.1 days. The global average value estimated by a Bayesian hierarchical model is shown. The value in each country is shown in **Extended Data Fig. 2d**. The posterior distribution (violin), 95% CI (line), and posterior mean (dot) are shown.

Although BA.1 spread worldwide earlier than BA.2, the lineage frequency of BA.2 increased and exceeded that of BA.1 since January 2022 in multiple countries, such as Philippines, India, Denmark, Singapore, Austria, and South Africa (**Fig. 1c** **and Extended Data Fig. 2**). To quantify the growth advantage of BA.2 in the human population, we constructed a Bayesian model representing the epidemic dynamics of SARS-CoV-2 lineages. This hierarchical model can estimate the global average of the relative effective reproduction numbers of viral lineages (**Fig. 1d**) as well as those in each country (**Extended Data Fig. 2**). The effective reproduction number of BA.2 is 1.40-fold higher than that of BA.1 on average in the world [95% confidence interval (CI), 1.29–1.52; **Fig. 1d**]. Furthermore, the effective reproduction number of BA.2 was even higher than that of BA.1.1, which spread more rapidly than BA.1 in several countries such as the UK and USA (**Fig. 1d** **and Extended Data Fig. 2d**). These results suggest that the BA.2 epidemic will more expand around the world, raising the importance of elucidating virological features of BA.2 in depth.

### Immune resistance of BA.2

Since the sequence of BA.2, particularly in S protein, is substantially different from that of BA.1 (**Fig. 1b** and **Fig. 2a**), it is reasonable to assume that the virological properties of BA.2, such as immune resistance and pathogenicity, are different from those of BA.1. To reveal the virological features of BA.2, we set out to perform neutralisation assay using pseudoviruses and the neutralising antibodies elicited by vaccination. Consistent with recent studies^4–11, 16–19^, BA.1 is highly resistant to the antisera elicited by mRNA-1273 and ChAdOx1 vaccines (**Fig. 2b,c**). Similar to BA.1, BA.2 was also highly resistant to the vaccine-induced antisera (**Fig. 2b,c**). Also, BA.2 was almost completely resistant to two therapeutic monoclonal antibodies, Casirivimab and Imdevimab, and was 35-fold more resistant to another therapeutic antibody, Sotrovimab, when compared to the ancestral D614G-bearing B.1.1 virus (**Fig. 2d**). Moreover, both BA.1 and BA.2 were highly resistant to the convalescent sera who had infected with early pandemic virus (before May 2020; **Fig. 2e**), Alpha (**Extended Data Fig. 3a**) and Delta (**Extended Data Fig. 3b**). These data suggest that, similar to BA.1, BA.2 is highly resistant to the antisera induced by vaccination and infection with other SARS-CoV-2 variants as well as three antiviral therapeutic antibodies.

**Fig. 2.**
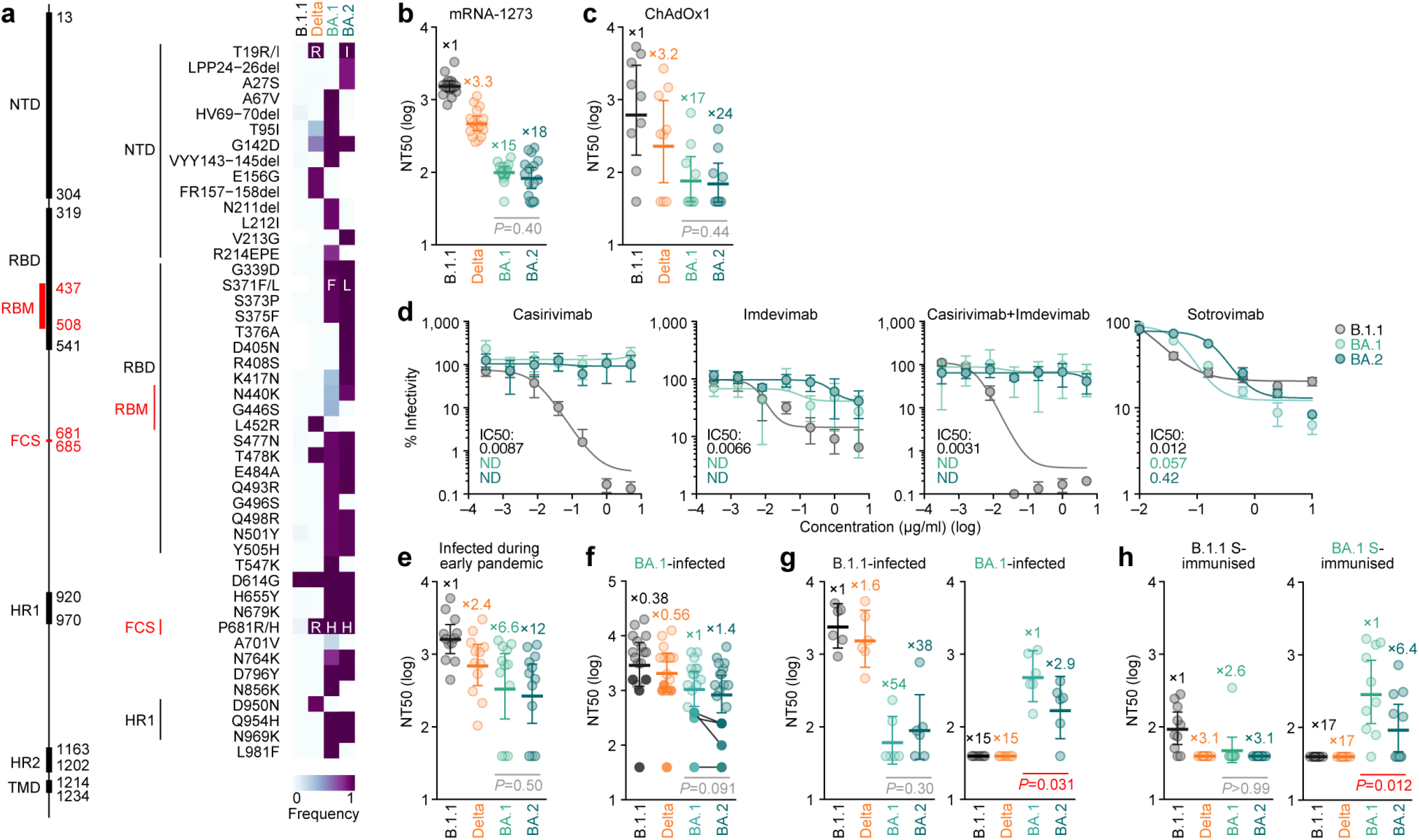
Immune resistance of BA.2. **a**, Amino acid substitutions in S. Left, primary structure and respective domains. Numbers indicate amino acid position. NTD, N-terminal domain; RBM, receptor-binding motif; HR, heptad repeat; TMD, transmembrane domain. Right, heatmap showing the frequency of amino acid substitutions. Substitutions detected in >10% sequences of any lineages are shown. **b–h**, Neutralisation assays. Neutralisation assays were performed with the pseudoviruses harbouring the S proteins of B.1.1 (the D614G-bearing ancestral virus), Delta, BA.1 and BA.2. Vaccinated sera [**b**, mRNA-1273 (16 donors); **c**, ChAdOx1 (9 donors)], monoclonal antibodies (**d**), convalescent sera of the individuals infected with early pandemic virus (until May 2020) (**e**, 12 donors), and BA.1 (**f**, 17 donors), convalescent sera of the hamsters infected B.1.1 (**g**, left; 6 hamsters) and BA.1 (**g**, right; 6 hamsters), and murine sera immunised with the cells expressing B.1.1 S (**h**, left; 10 mice) and BA.1 S (**h**, right; 10 mice) were used. In **b,c,e–h**, assay of each serum was performed in triplicate to determine 50% neutralisation titre (NT50). Each dot represents each NT50 value, and geometric mean and 95% CI are shown. Number indicates the fold change of resistance versus each antigenic variant. Statistically significant differences between BA.1 and BA.2 were determined by two-sided Wilcoxon signed-rank test. Information of vaccinated/convalescent donors are summarized in **Supplementary Tables 1 and 2**. In **f**, the dots of not-fully vaccinated 4 donors are filled. In **d**, assay of each concentration was performed in triplicate, and data are the average ± s.d. IC50, 50% inhibitory concentration; ND, not determined.

We then tested the 17 sera infected with BA.1: 13 convalescents were fully vaccinated (2 shots), 1 convalescent was 1-dose vaccinated, and 3 convalescents were not vaccinated. BA.1 convalescent sera exhibited the strongest antiviral effect against BA.1 (**Fig. 2f**). Although BA.2 was 1.4-fold more resistant to the BA.1-infected sera than BA.1, there was no statistical difference (**Fig. 2f**; *P*=0.091 by Wilcoxon signed-rank test). Importantly, the BA.1 convalescent sera with full vaccination exhibited significantly stronger antiviral effects against all variants tested than unvaccinated or 1-dose vaccinated convalescents (**Extended Data Fig. 3c**).

To address the possibility that the BA.1-induced humoral immunity is less effective against BA.2, we used the convalescent sera obtained from infected hamsters at 16 days postinfection (d.p.i.). Similar to the results of convalescent human sera (**Fig. 2e** and **Extended Data Fig. 2b**), both BA.1 and BA.2 exhibited pronounced resistances against B.1.1- and Delta-infected convalescent hamster sera (**Fig. 2g** and **Extended Data Fig. 3d**). Interestingly, BA.2 was significantly (2.9-fold) more resistant to BA.1-infected convalescent hamster sera than BA.1 (**Fig. 2g**). To further verify the resistance of BA.2 against BA.1-induced immunity, mice were immunised with the cells expressing the S proteins of ancestral B.1.1 and BA.1 and obtained murine antisera. Again, the neutralisation assay using murine sera showed that BA.2 is more significantly (6.4-fold) resistant to the BA.1 S-immunised sera than BA.1 (**Fig. 2h**). These findings suggest that BA.1-induced humoral immunity is less effective against BA.2.

### Virological features of BA.2 in vitro

To investigate the virological characteristics of BA.2, we generated chimeric recombinant SARS-CoV-2 that expresses GFP and harbours the *S* gene of ancestral B.1.1, Delta, BA.1 and BA.2 by reverse genetics (**Extended Data Fig. 4**)^21^. Although the growth of BA.1 and BA.2 was comparable in VeroE6/TMPRSS2 cells, BA.2 was more replicative than BA.1 in Calu-3 cells and primary human nasal epithelial cells (**Fig. 3a**). Notably, the morphology of infected cells was different; BA.2 formed significantly (1.52-fold) larger syncytia than BA.1 (**Fig. 3b** and **Extended Data Fig. 5a**). Whereas the plaque size in VeroE6/TMPRSS2 cells infected with BA.1 and BA.2 was significantly smaller than those of cells infected with B.1.1, the plaques formed by BA.2 infection are significantly (1.27-fold) larger than those by BA.1 infection (**Fig. 3c** and **Extended Data Fig. 5b**). Moreover, the coculture of S-expressing cells with HEK293-ACE2/TMPRSS2 cells showed that BA.2 S induces significantly (2.9-fold) larger multinuclear syncytia formation when compared to BA.1 S (**Extended Data Fig. 5c**). These data suggest that BA.2 is more fusogenic than BA.1. To further address this possibility, we analysed the fusogenicity of the S proteins of BA.2 S by a cell-based fusion assay^12, 22, 23^. The expression level of BA.2 S on the cell surface was significantly lower than that of BA.1 S (**Extended Data Fig. 6a**). Nevertheless, our fusion assay using VeroE6/TMPRSS2 cells and Calu-3 cells showed that BA.2 S is significantly more fusogenic than BA.1 S (**Fig. 3d**). We then analysed the binding affinity of BA.2 S receptor binding domain (RBD) to ACE2 by an yeast surface display assay^16, 22, 24^. Although the binding affinity of BA.1 S RBD to ACE2 is controversial^11, 15–17, 25, 26^, our yeast surface display showed that the binding affinity of the RBD of BA.1 and BA.2 is comparable (**Extended Data Fig. 6b**).

**Fig. 3.**
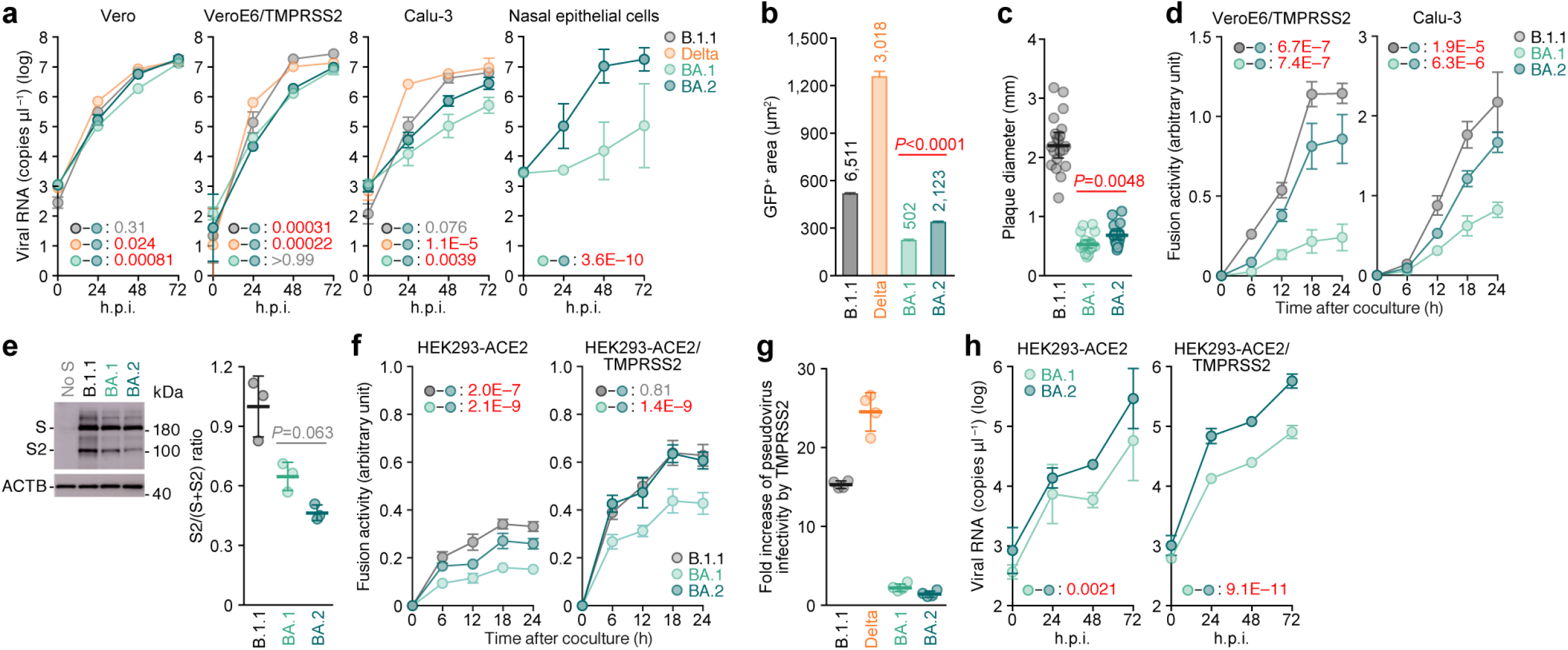
Virological features of BA.2 in vitro. **a,h**, Growth kinetics of SARS-CoV-2 variants. **b**, Fluorescence microscopy. GFP area of infected VeroE6/TMPRSS2 cells at 48 h.p.i were measured. Numbers in the panel indicate the number of GFP-positive cells counted. **c**, Plaque assay. Diameter of plaques (20 plaques per virus) are summarized. **d,f**, S-based fusion assay. The fusion activity (arbitrary units) is shown. **e**, Western blot. Left, representative blots of S-expressing cells. ACTB is an internal control. Right, the ratio of S2 to the full-length S plus S2 proteins. **g**, Fold increase of pseudovirus infectivity by TMPRSS2 expression. Assays were performed in quadruplicate (**a,g,h**), octuplicate (**a**, most left) or triplicate (**d–f**) and data are the average ± s.d. Each dot indicates the result from an individual plaque (**c**) and an individual replicate (**e,g**). In **b** and **c**, raw data are shown in **Extended Data Fig. 5 and 6**. Statistically significant differences between BA.2 and other variants through timepoints were determined by multiple regression (**a,d,f,h**). Familywise error rates (FWERs) calculated using the Holm method are indicated in the figures. Statistically significant differences between BA.1 and BA.2 were determined by two-sided Mann–Whitney *U*-tests (**b,d**) or two-sided paired Student’s *t*-tests (**e**).

Because we have proposed that the SARS-CoV-2 S-mediated fusogenicity is closely associated with the efficacy of S1/S2 cleavage^12, 23^, we hypothesized that BA.2 S is more efficiently cleaved than BA.1 S. However, an western blotting analysis showed that BA.2 S is less cleaved than BA.1 S (**Fig. 3e**), suggesting that BA.2 S exhibits a higher fusogenicity independently of S1/S2 cleavage.

We have recently revealed that BA.1 poorly utilizes TMPRSS2 for the infection^11^. To analyse the TMPRSS2 usage by BA.2 S, we performed cell-based fusion assay using 293-ACE2 cells with or without TMPRSS2 expression. We verified that 293-ACE2 cells do not express endogenous TMPRSS2 on the cell surface (**Extended Data Fig. 6c**). As shown in **Fig. 3f**, the fusogenicity of BA.2 S was significantly higher in both cell lines than that of BA.1. However, although BA.2 S was less fusogenic than B.1.1 S in 293-ACE2 cells, the fusogenicity of BA.2 S and B.1.1 S was comparable in 293-ACE2/TMPRSS2 cells (**Fig. 3f**). These results suggest that the relatively higher fusogenicity of BA.2 is dependent on TMPRSS2 expression on the surface of target cells. To further assess whether the TMPRSS2-dependent infection enhancement was also observed in cell-free virus, we inoculated pseudoviruses into 293-ACE2 cells and 293-ACE2/TMPRSS2 cells. Although the infectivity of B.1.1 and Delta was 15.3-fold and 24.6-fold increased by TMPRSS2 expression, respectively, the TMPRSS2 expression on the target cells did not affect the infectivity of both BA.1 and BA.2 (**Fig. 3g**). These results suggest that TMPRSS2 does not affect the infectivity of cell-free BA.2 virus. However, although the growth of BA.2 and BA.1 was comparable in 293-ACE2 cells, BA.2 was more replicative than BA.1 in 293-ACE2/TMPRSS2 cells (**Fig. 3h**). Overall, our data suggest that BA.2 is more fusogenic and replicative than BA.1 in a TMPRSS2-dependent fashion.

### Virological features of BA.2 in vivo

To investigate the dynamics of viral replication of BA.2 *in vivo*, we conducted hamster infection experiments. Consistent with our recent study^12^, B.1.1-infected hamsters exhibited decreased body weight and respiratory disorders that are reflected by two surrogate markers for bronchoconstriction or airway obstruction, enhanced pause (Penh) and the ratio of time to peak expiratory follow relative to the total expiratory time (Rpef), and subcutaneous oxygen saturation (SpO_2_), whereas BA.1-infected hamsters exhibited no or weak disorders (**Fig. 4a**). Notably, all parameters routinely measured, including body weight, Penh, Rpef and SpO_2_, of BA.2-infected hamsters were significantly different from uninfected and BA.1-infected hamsters, and these values were comparable to those of B.1.1-infected hamsters (**Fig. 4a**). These data suggest that BA.2 is more pathogenic than BA.1.

**Fig. 4.**
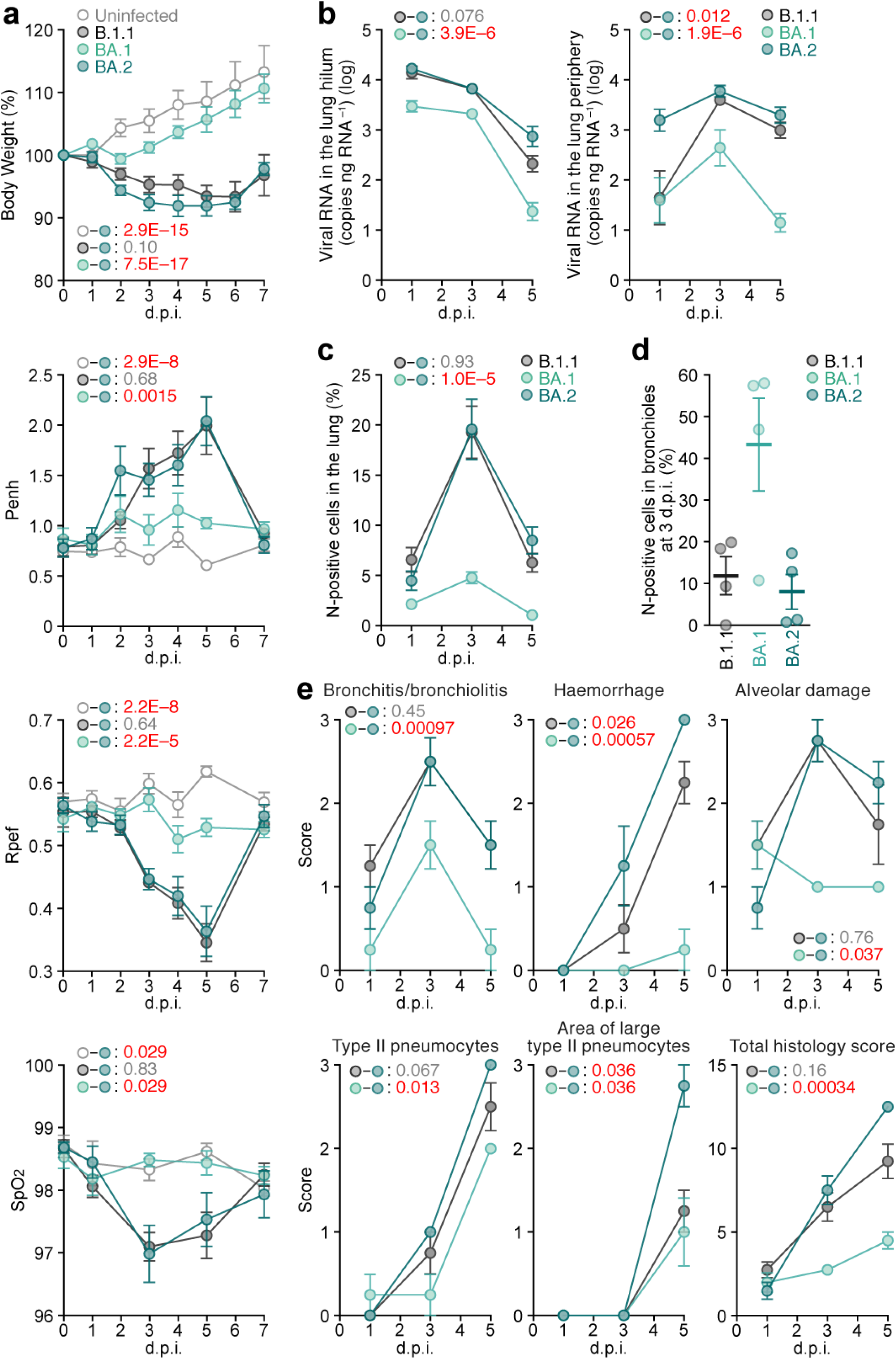
Virological features of BA.2 in vivo. Syrian hamsters were intranasally inoculated with B.1.1, BA.1 and BA.2. **a**, Body weight, Penh, Rpef, SpO_2_ values routinely measured. Hamsters of the same age were intranasally inoculated with PBS (uninfected). **b**, Viral RNA load in the lung hilum (left) and periphery (right). **c,d**, Percentage of N-positive cells in the whole lobes of lung (**c**) and bronchiole in the frontal/upper lobe of lung (**d**) measured by IHC. **e**, Histopathological scoring of lung lesions. Representative pathological features are shown in our previous studies^12, 23^. Data are the average (**a**, 6 hamsters per each group; **b–e**, 4 hamsters per each group) ± s.e.m. In **a–c,e**, statistically significant differences between BA.2 and other variants or uninfected hamsters through timepoints were determined by multiple regression. The data at 0 d.p.i. was excluded from the analyses. FWERs calculated using the Holm method are indicated in the figures. In **d**, each dot indicates the result from an individual hamster. In **c–e**, raw data are shown in **Extended Data Fig. 7 and 8**.

To analyse viral spread in the respiratory organs of infected hamsters, viral RNA load and nucleocapside (N) expression were assessed by RT-qPCR of viral RNA and immunohistochemistry (IHC), respectively. As shown in **Fig. 4b**, viral RNA loads in the two lung regions, hilum and periphery, of BA.2-infected hamsters were significantly higher than those of BA.1-infected hamsters. In the lung periphery, the viral RNA load of BA.2 was significantly higher than that of B.1.1, and the viral RNA load of BA.2 at 1 d.p.i. was 11-fold and 9.3-fold higher than those of B.1.1 and BA.1 at the same timepoint, respectively (**Fig. 4b**). To address the possibility that BA.2 more efficiently spreads than BA.1, we investigated the positivity for N protein in the trachea and lung area close to the hilum. At 1 d.p.i., N protein was detectable in the lower tracheal epithelium in all infected hamsters, and particularly, was clearly detectable in the middle part of trachea in BA.2-infected hamsters (**Extended Data Fig. 7a**). The positivity of N protein was observed in both bronchial and bronchiolar epithelium in all infected lungs (**Extended Data Fig. 7b**). Notably, alveolar positivities were observed in B.1.1- and BA.2-infected lungs but not in BA.1-infected lungs (**Extended Data Fig. 7b**). Morphometry showed that the percentage of N-positive cells in BA.2-infected lungs is significantly higher than that of BA.1-infected lungs and peaked at 3 d.p.i. (**Fig. 4c** and **Extended Data Fig. 7c**). On the other hand, at 3 d.p.i., the percentage of N-positive cells in the bronchus/bronchioles of BA.2-infected hamsters was 5.4-fold lower than that of BA.1-infected hamsters (**Fig. 4d**). At 5 d.p.i., N protein was almost disappeared in BA.1-infected lungs, whereas alveolar staining was still detectable in B.1.1- and BA.2-infected lungs (**Fig. 4c** and **Extended Data Fig. 7c**). These data suggest that BA.2 is more rapidly and efficiently spread in the lung tissues than BA.1.

### Pathogenicity of BA.2

To investigate the pathogenicity of BA.2, the right lungs of infected hamsters were collected at 1, 3, and 5 d.p.i. and used them for haematoxylin and eosin (H&E) staining and histopathological analysis^12, 23^. All histopathological parameters including bronchitis/bronchiolitis, haemorrhage, alveolar damage, and the levels of type II pneumocytes, of BA.2-infected hamsters were significantly higher than those in BA.1 (**Fig 4e** and **Extended Data Fig. 8a**). The score indicating haemorrhage including congestive edema of BA.2 was significantly higher than that of B.1.1 (**Fig. 4e**). As shown in our previous studies^12, 23^, hyperplastic large type II pneumocytes suggesting the severity of inflammation were observed in all infected hamsters at 5 d.p.i., and particularly, the area of large type II pneumocytes in BA.2-infected hamsters was significantly larger than those in B.1.1- and BA.1-infected hamsters (**Fig. 4e**). Total histology score of BA.2 was significantly higher than that of BA.1 (**Fig. 4e**). Furthermore, in the BA.2- and B.1.1-infected lungs, the inflammation with type II alveolar pneumocyte hyperplasia was found in each lobe especially frontal/upper and accessary lobes (**Extended Data Fig. 8b**).

## Discussion

Although BA.2 is considered as an Omicron variant, its genomic sequence is heavily different from BA.1, which suggests that the virological characteristics of BA.2 is different from that of BA.1. Here, we elucidated the virological characteristics of BA.2, such as its higher effective reproduction number, higher fusogenicity, higher pathogenicity when compared to BA.1. Moreover, we demonstrated that BA.2 is resistant to the BA.1-induced humoral immunity. Our data indicate that BA.2 is virologically different from BA.1 and raise a proposal that BA.2 should be given a letter of the Greek alphabet and be distinguished from BA.1, a commonly recognized Omicron variant.

We showed evidence suggesting that BA.2 is virologically different and distinguishable from BA.1. First, by using the two different types of antisera that were obtained from experimental animals, convalescent hamsters infected with BA.1 and mice immunized with the BA.1 S protein, we showed that the resistance of BA.2 to the BA.1-induced humoral immunity. Our results indicate that the antigenicity of BA.2 is different from BA.1. Although similar tendency was observed in human samples as well, a statistically significant difference was not observed probably because a relatively lower number of vaccine-naïve individuals who were infected with BA.1 (3 unvaccinated donors and a 1-dose vaccinated donor) were tested in the present study. The effect of BA.1-induced humoral immunity against BA.2 in humans should deeply be verified in the future investigation.

Second, the higher fusogenicity of BA.2 S is a pronounced characteristics in in vitro experiments. We have demonstrated that Delta S is highly fusogenic than BA.1 S and B.1.1 S, and we considered that the higher fusogenicity is attributed to the higher efficacy of S cleavage^11, 12, 23, 27^. However, BA.2 S exhibited higher fusogenicity than BA.1 S without the increase of S cleavage efficacy. In recent studies^12, 23^, we have proposed that the fusogenicity of SARS-CoV-2 variant is closely related to its pathogenicity. This hypothesis is further supported by the observations in BA.2 in the present study. However, unlike Delta, the higher fusogenicity of BA.2 appears to be not attributed to the higher efficacy of S cleavage^11, 12, 23, 27^. Moreover, although TMPRSS2 increased the efficacies of both cell-cell fusion^23^ and cell-free infection mediated by B.1.1 S and Delta S, TMPRSS2 increased the efficacy of BA.2 S-mediated cell-cell fusion but did not affect that of BA.2 S-mediated cell-free infection. These observations suggest that TMPRSS2 contributes to the cell-cell fusion and cell-free infection mediated by BA.2 S with different mechanisms of action.

Third, it would be most critical for global health that BA.2 exhibits higher pathogenicity than BA.1. Although clinical researches on the BA.2 pathogenicity are needed, our investigations using a hamster model showed that the pathogenicity of BA.2 is similar to that of an ancestral B.1.1 and higher than that of BA.1. More importantly, the viral RNA load in the lung periphery and histopathological disorders of BA.2 were more severe than those of BA.1 and even B.1.1. Together with a higher effective reproduction number and pronounced immune resistance of BA.2, it is evident that the spread of BA.2 can be a serious issue for global health in the near future.

In summary, our data suggest the possibility that BA.2 would be the most concerned variant to global health. Currently, both BA.2 and BA.1 are recognised together as Omicron and these are almost undistinguishable. Based on our findings, we propose that BA.2 should be recognised as a unique variant of concern, and this SARS-CoV-2 variant should be monitored in depth.

## Methods

### Ethics statement

All experiments with hamsters were performed in accordance with the Science Council of Japan’s Guidelines for the Proper Conduct of Animal Experiments. The protocols were approved by the Institutional Animal Care and Use Committee of National University Corporation Hokkaido University (approval ID: 20-0123 and 20-0060). All experiments with mice were also performed in accordance with the Science Council of Japan’s Guidelines for the Proper Conduct of Animal Experiments. The protocols were approved by the Institutional Animal Experiment Committee of The Institute of Medical Science, The University of Tokyo (approval ID: PA21-39). All protocols involving specimens from human subjects recruited at Kyoto University, Kuramochi Clinic Interpark and Chiba University were reviewed and approved by the Institutional Review Boards of Kyoto University (approval ID: G1309), Kuramochi Clinic Interpark (approval ID: G2021-004) and Chiba University (approval ID: HS202103-03). All human subjects provided written informed consent. All protocols for the use of human specimens were reviewed and approved by the Institutional Review Boards of The Institute of Medical Science, The University of Tokyo (approval IDs: 2021-1-0416 and 2021-18-0617), Kyoto University (approval ID: G0697), Kumamoto University (approval IDs: 2066 and 2074), and University of Miyazaki (approval ID: O-1021).

### Human sera collection

Vaccine sera were collected from sixteen vaccinees four weeks after the second vaccination with mRNA-1273 (Moderna) (average age: 27, range: 20-47, 38% male). The sera obtained from nine vaccinees 10-25 d after the second vaccination with ChAdOx1 (Oxford-AstraZeneca) (average age: 45, range: 35-54, 67% male) were purchased from BioIVT. The detail of vaccine sera is summarized in **Supplementary Table 1**.

Convalescent sera were collected from vaccine-naive individuals who had infected with Alpha variant (n=8; average age: 41, range: 21-57, 63% male) and Delta variant (n=15; average age: 51, range: 22-67, 80% male). Convalescent sera of BA.1-infected individuals (n=17; average age: 39, range: 20-65, 47% male, 76% received the second vaccination) were also collected. To determine the SARS-CoV-2 variants infected, salivas were collected from COVID-19 patients during onset and RNA was extracted using a QIAamp viral RNA mini kit (Qiagen, Cat# 52906) according to the manufacturer’s protocol. To determine Alpha and Delta variants, viral genome sequencing was performed as previously described^11^. For the detail, see “Viral genome sequencing” section below. To determine the BA.1 variant, mutation-targeting RT-qPCR was performed. To detect the S E484A substitution (common in all Omicron variants including BA.1 and BA.2), a set of primer/probe E484A (SARS-CoV-2) (Takara, Cat# RC322A) was used. To detect the S R214EPE insertion (specific for B.1.1.529 and BA.1, undetectable in BA.2), a hand-made protocol was used with the following primers and probe: Omi_ins214s-F1, TTC TAA GCA CAC GCC TAT TAT AGT GC; Omi_ins214s-R1, TAA AGC CGA AAA ACC CTG AGG; and Omi_ins214s, FAM-TGA GCC AGA AGA TC-MGB. The twelve convalescent sera during early pandemic (until May 2020) (average age: 71, range: 52-92, 8% male) were purchased from RayBiotech. Sera were inactivated at 56°C for 30 min and stored at –80°C until use. The detail of convalescent sera is summarized in **Supplementary Table 2**.

### Viral genome sequencing

Viral genome sequencing was performed as previously described^11, 12, 22, 23^ with some modifications. Briefly, the sequences of the working viruses were verified by viral RNA-sequencing analysis. Viral RNA was extracted using QIAamp viral RNA mini kit (Qiagen, Cat# 52906). The sequencing library for total RNA-sequencing was prepared using NEB Next Ultra RNA Library Prep Kit for Illumina (New England Biolabs, Cat# E7530). Paired-end, 76-bp sequencing was performed using MiSeq (Illumina) with MiSeq reagent kit v3 (Illumina, Cat# MS-102-3001). Sequencing reads were trimmed using fastp v0.21.0^28^ and subsequently mapped to the viral genome sequences of a lineage A isolate (strain WK-521; GISAID ID: EPI_ISL_408667)^29^ using BWA-MEM v0.7.17^30^. Variant calling, filtering, and annotation were performed using SAMtools v1.9^31^ and snpEff v5.0e^32^.

### Phylogenetic and comparative genome analyses

To construct a maximum likelihood tree of the Omicron lineages (BA.1, BA.1.1, BA.2, and BA.3) sampled from South Africa, the genome sequence data of SARS-CoV-2 and its metadata were downloaded from the GISAID database (https://www.gisaid.org/) on January 26, 2022. We excluded the data of viral strains with the following features from the analysis: i) lacking the collection date information; ii) sampled from animals other than humans; iii) having the flag of low coverage sequencing; or iv) having >2% of undetermined nucleotide characters. All the BA.2 and BA.3 sequences and 200 randomly sampled BA.1 (including 20 BA.1.1) sequences were used for the tree construction in addition to an outgroup sequence, EPI_ISL_466615, the oldest isolate of B.1.1 in the UK. The viral genome sequences were mapped to the reference sequence of Wuhan-Hu-1 (GenBank accession no.: NC_045512.2) using minimap2 v2.17^33^ and subsequently converted to the multiple sequence alignment according to the GISAID phylogenetic analysis pipeline (https://github.com/roblanf/sarscov2phylo). The alignment sites corresponding to the 1–265 and 29674–29903 positions in the reference genome were masked (i.e., converted to NNN). The alignment sites with >50% sequences having a gap or undetermined/ambiguous nucleotide were trimmed using trimAl v1.2^34^. The phylogenetic tree construction was performed by a two-step protocol: i) The first tree was constructed; ii) tips with a longer external branch (Z score > 4) were removed from the dataset; iii) and the final tree was constructed. The tree reconstruction was performed by RAxML v8.2.12^35^ under the GTRCAT substitution model. Node support value was calculated by 100-times of bootstrap analysis.

We performed phylodynamics analysis of Omicron lineages (BA.1, BA.1.1, BA.2 and BA.3) sampled from South Africa as below. The SARS-CoV-2 genome sequence dataset used above was split into each Omicron lineage. As an outgroup sequence, the oldest BA.2 (GISAID ID: EPI_ISL_8128463) was added to the BA.1 and BA.3 datasets, and the oldest BA.3 (GISAID ID: EPI_ISL_8616600) was added to the BA.2 dataset. The multiple sequence alignment was constructed as the procedures described above. A time-calibrated tree of each lineage was constructed by BEAST2 v.2.6.6^36^. The HKY model^37^ with the four categories of discrete gamma rate variation was selected as a nucleotide substitution model. A relaxed molecular clock modelled by a log-normal distribution was selected. The exponential growth coalescent model was used. For BA.1, BA.2, and BA.3 datasets, nineteen, four, and three independent chains of Markov chain Monte Carlo (MCMC) were respectively run with 2,000,000 warmup and 18,000,000 sampling iterations. We confirmed that the effective sampling sizes for all parameters were greater than 200, indicating the MCMC runs were successfully convergent. The maximum credible trees with common ancestor heights are shown in **Extended Data Fig. 1**.

The number of amino acid differences (including nonsynonymous substitutions, insertions, and deletions) between SARS-CoV-2 lineages was defined as below. Information on amino acid differences of each viral strain compared with the reference sequence of Wuhan-Hu-1 (GenBank accession no.: NC_045512.2) was extracted from the GISAID metadata (downloaded on January 26, 2022). In each viral lineage, the amino acid differences that were present in >10% sequences were extracted and subsequently counted. For the comparison of BA.1 and BA.2, the set of the symmetric difference of the amino acid differences compared with the reference was determined and subsequently counted.

### Modelling the epidemic dynamics of SARS-CoV-2 lineages

To quantify the spread speed of each SARS-CoV-2 lineage in the human population, we estimated the relative effective reproduction number of each viral lineage according to the epidemic dynamics calculated on the basis of viral genomic surveillance data. The data were downloaded from the GISAID database (https://www.gisaid.org/) on February 1, 2022. We excluded the data of viral strains with the following features from the analysis: i) lacking the collection date information; ii) sampled from animals other than humans; or iii) sampled by quarantine. We analysed the datasets of the eleven countries with >100 BA.2 sequences (Austria, Denmark, Germany, India, Israel, Philippines, Singapore, South Africa, Sweden, the UK, and the USA) (**Fig. 1** **and Extended Data Fig. 2**). The dynamics of BA.1, BA.1.1, BA.2, and Delta (B.1.617.2 and AY lineages) in each country from October 1, 2021, to January 25, 2022, were analysed. The number of viral sequences of each viral lineage collected on each day in each country was counted. Finally, we constructed the L (lineage) × C (country) × T (time) = 4 × 11 × 117 shaped array, which comprises the count of each viral lineage in each country in each day. This array was used as input data for the statistical model described below.

We constructed a Bayesian hierarchical model to represent the relative lineage growth dynamics with a multinomial logistic regression. The mathematical theory underlying the model is described in detail elsewhere^38, 39^, and this model is basically similar to the one used in our previous study^12^. In the present study, we incorporated a hierarchical structure into the slope parameter over time, which enable us to estimate the global average of the relative effective reproduction number of each viral lineage as well as that in each country. Arrays in the model index over one or more indices: L = 4 viral lineages *l*; C = 11 countries *c*; and T = 117 days *t*. The model is:

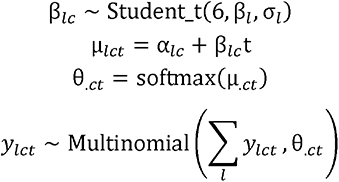

The explanatory variable is time *t*, and the outcome variable is *y_lct_*, which represents the count of viral lineage *l* in country *c* at time *t*. The slope parameter of lineage *I* in country *c*, *β_lc_*, is generated from a student’s t distribution with the hyper parameters of the mean *β_l_* and the standard deviation *σ_l_*. As the distribution generating *β_lc_*, we used a student’s t distribution with six degrees of freedom instead of a normal distribution to reduce the effects of outlier values of *β_lc_*. In the model, the linear estimator μ_.*ct*_, consisting of the intercept α*_.c_* and the slope β*_.c_*, is converted to the simplex θ*_.ct_*, which represents the probability of occurrence of each viral lineage at time *t* in country *c*, by the softmax link function defined as:

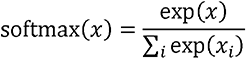

*y_lct_* is generated from θ*_.ct_* and the total count of all lineages at *t* in country *c* according to a multinomial distribution.

The relative effective reproduction number of each viral lineage in each county (*r_lc_*) was calculated according to the slope parameter *β_lc_* as:

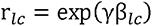

where γ is the average viral generation time (2.1 day) (http://sonorouschocolate.com/covid19/index.php?title=Estimating_Generation_Time_Of_Omicron). Similarly, the global average of the relative effective reproduction number of each viral lineage was calculated according to the slope hyper parameter *β_l_* as:

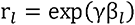

For the parameter estimation, the intercept and slope parameters of the BA.1 variant were fixed at 0. Consequently, the relative effective reproduction number of BA.1 was fixed at 1, and those of the respective lineages were estimated relative to that of BA.1.

Parameter estimation was performed by MCMC implemented in CmdStan v2.28.1 (https://mc-stan.org) with cmdstanr v0.4.0 (https://mc-stan.org/cmdstanr/). Noninformative priors were set for all parameters. Four independent MCMC chains were run with 1,000 and 2,000 steps in the warmup and sampling iterations, respectively. We confirmed that all estimated parameters had <1.01 R-hat convergence diagnostic and >200 effective sampling size values, indicating that the MCMC runs were successfully convergent. The fitted model closely recapitulated the observed viral lineage dynamics in each country (**Extended Data Fig. 2c**). The analyses above were performed in R v4.1.2 (https://www.r-project.org/).

### Cell culture

HEK293 cells (a human embryonic kidney cell line; ATCC CRL-1573) and HEK293-ACE2 cells [HEK293 cells (ATCC CRL-1573) stably expressing human ACE2]^22^ were maintained in DMEM (high glucose) (Sigma-Aldrich, Cat# 6429-500ML) containing 10% FBS, 1 µg/ml puromycin (InvivoGen, Cat# ant-pr-1) and 1% PS. HEK293-ACE2/TMPRSS2 cells [HEK293 cells (ATCC CRL-1573) stably expressing human ACE2 and TMPRSS2]^22^ were maintained in Dulbecco’s modified Eagle’s medium (DMEM) (high glucose) (Wako, Cat# 044-29765) containing 10% foetal bovine serum (FBS) and 1% penicillin-streptomycin (PS). HEK293-C34 cells, *IFNAR1* KO HEK293 cells expressing human ACE2 and TMPRSS2 by doxycycline treatment^21^, were maintained in Dulbecco’s modified Eagle’s medium (high glucose) (Sigma-Aldrich, Cat# R8758-500ML) containing 10% FBS, 10 μg/ml blasticidin (InvivoGen, Cat# ant-bl-1) and 1% PS. Vero cells [an African green monkey (*Chlorocebus sabaeus*) kidney cell line; JCRB0111] were maintained in Eagle’s minimum essential medium (EMEM) (Wako, Cat# 051-07615) containing 10% FBS and 1% PS. VeroE6/TMPRSS2 cells (VeroE6 cells stably expressing human TMPRSS2; JCRB1819)^29^ were maintained in DMEM (low glucose) (Wako, Cat# 041-29775) containing 10% FBS, G418 (1 mg/ml; Nacalai Tesque, Cat# G8168-10ML) and 1% PS. Calu-3 cells (a human lung epithelial cell line; ATCC HTB-55) were maintained in EMEM (Sigma-Aldrich, Cat# M4655-500ML) containing 20% FBS and 1% PS. Calu-3/DSP_1-7_ cells [Calu-3 cells (ATCC HTB-55) stably expressing DSP_1-7_]^40^ were maintained in EMEM (Wako, Cat# 056-08385) supplemented with 20% FBS and 1% PS. B16F10 cells (a mouse melanoma cell line; RCB2630) were maintained in DMEM (high glucose) (Sigma-Aldrich, Cat# D6429-500ML) containing 10% FBS and 1% PS. Expi293 cells (Thermo Fisher Scientific, Cat# A14527) were maintained in Expi293 expression medium (Thermo Fisher Scientific, Cat# A1435101). Primary human nasal epithelial cells (Cat# EP02, Batch# MP0010) were purchased from Epithelix and maintained according to the manufacturer’s procedure.

### Plasmid construction

Plasmids expressing the SARS-CoV-2 S proteins of B.1.1 (the parental D614G-bearing variant), Alpha (B.1.1.7), Delta (B.1.617.2) and BA.1 variants were prepared in our previous studies^11, 12, 22, 23, 41^. Plasmids expressing the codon-optimised S proteins of BA.2 and a BA.2 derivative that loses its cytoplasmic tail were generated by site-directed overlap extension PCR using the primers listed in **Supplementary Table 3**. The resulting PCR fragment was digested with KpnI and NotI and inserted into the corresponding site of the pCAGGS vector. To construct the plasmids expressing anti-SARS-CoV-2 monoclonal antibodies (Casirivimab, Imdevimab or Sotrovimab), the sequences of the variable regions of these antibodies were obtained from KEGG Drug Database (https://www.genome.jp/kegg/drug/) and were artificially synthesized (Fasmac). The obtained coding sequences of the variable regions of the heavy and light chains were cloned into the pCAGGS vector containing the sequences of the human immunoglobulin 1 and kappa constant region [kindly provided by Dr. Hisashi Arase (Osaka University, Japan)]. Nucleotide sequences were determined by DNA sequencing services (Eurofins), and the sequence data were analyzed by Sequencher v5.1 software (Gene Codes Corporation).

### Preparation of monoclonal antibodies

Casirivimab, Imdevimab and Sotrovimab were prepared as previously described^11, 42^. Briefly, the pCAGGS vectors containing the sequences encoding the immunoglobulin heavy and light chains were cotransfected into HEK293T cells using PEI Max (Polysciences, Cat# 24765-1). At 48 h posttransfection, the cell culture supernatants were harvested, and the antibodies were purified using NAb protein A plus spin kit (Thermo Fisher Scientific, Cat# 89948) according to the manufacturer’s protocol.

### Preparation of mouse sera

The SARS-CoV-2 S-immunised mouse sera were prepared as previously described^42^. To prepare the immunogen, B16F10 cells (2,500,000 cells) were transfected with 5 μg S expression plasmid by PEI Max (Polysciences, Cat# 24765-1) according to the manufacturer’s protocol. Two days posttransfection, the transfected cells were washed twice with PBS, and then the cell pellets were stored at –80°C (10,000,000 cells per stock). The expression of transfected S protein was verified by flow cytometry and western blot. BALB/c mice (female, 7 weeks old) were purchased from Japan SLC Inc. (Shizuoka, Japan). The mice were maintained under specific pathogen-free conditions. For the immunisation, mice were subcutaneously immunized with the freeze-thawed S-expressing B16F10 cells in complete Freund’s adjuvant (50%) (Sigma-Aldrich, Cat# F5881). Three weeks after immunisation, blood was collected in BD Microtainer blood collection tubes (BD Biosciences, Cat# 365967) and sera were collected by centrifugation.

### Neutralisation assay

Pseudoviruses were prepared as previously described^11, 23, 24, 41, 43^. Briefly, lentivirus (HIV-1)-based, luciferase-expressing reporter viruses were pseudotyped with the SARS-CoV-2 spikes. HEK293T cells (1,000,000 cells) were cotransfected with 1 μg psPAX2-IN/HiBiT^44^, 1 μg pWPI-Luc2^44^, and 500 ng plasmids expressing parental S or its derivatives using PEI Max (Polysciences, Cat# 24765-1) according to the manufacturer’s protocol. Two days posttransfection, the culture supernatants were harvested and centrifuged. The pseudoviruses were stored at –80°C until use.

For the neutralisation assay, the SARS-CoV-2 S pseudoviruses (counting ∼20,000 relative light units) were incubated with serially diluted (40-fold to 29,160-fold dilution at the final concentration) heat-inactivated sera or monoclonal antibodies (Casirivimab, Imdevimab or Sotrovimab) at 37°C for 1 h. Pseudoviruses without sera were included as controls. Then, an 80 μl mixture of pseudovirus and serum/antibody was added to HOS-ACE2/TMPRSS2 cells (10,000 cells/50 μl) in a 96-well white plate. At 2 d.p.i., the infected cells were lysed with a One-Glo luciferase assay system (Promega, Cat# E6130) or a Bright-Glo™ Luciferase Assay System (Promega, Cat# E2650), and the luminescent signal was measured using a GloMax explorer multimode microplate reader 3500 (Promega) or CentroXS3 (Berthhold Technologies). The assay of each serum was performed in triplicate, and the 50% neutralisation titre (NT50) was calculated using Prism 9 (GraphPad Software).

### SARS-CoV-2 reverse genetics

Recombinant SARS-CoV-2 was generated by circular polymerase extension reaction (CPER) as previously described^21–23^. In brief, 9 DNA fragments encoding the partial genome of SARS-CoV-2 (strain WK-521, PANGO lineage A; GISAID ID: EPI_ISL_408667)^29^ were prepared by PCR using PrimeSTAR GXL DNA polymerase (Takara, Cat# R050A). A linker fragment encoding hepatitis delta virus ribozyme, bovine growth hormone poly A signal and cytomegalovirus promoter was also prepared by PCR. The corresponding SARS-CoV-2 genomic region and the PCR templates and primers used for this procedure are summarised in **Supplementary Table 4**. The 10 obtained DNA fragments were mixed and used for CPER^21^. To prepare GFP-expressing replication-competent recombinant SARS-CoV-2, we used fragment 9, in which the *GFP* gene was inserted in the *ORF7a* frame, instead of the authentic F9 fragment (see **Extended Data Fig. 4** and **Supplementary Table 4**)^21^.

To produce recombinant SARS-CoV-2, the CPER products were transfected into HEK293-C34 cells using TransIT-LT1 (Takara, Cat# MIR2300) according to the manufacturer’s protocol. At one day posttransfection, the culture medium was replaced with Dulbecco’s modified Eagle’s medium (high glucose) (Sigma-Aldrich, Cat# R8758-500ML) containing 2% FCS, 1% PS and doxycycline (1 μg/ml; Takara, Cat# 1311N). At six days posttransfection, the culture medium was harvested and centrifuged, and the supernatants were collected as the seed virus. To remove the CPER products (i.e., SARS-CoV-2-related DNA), 1 ml of the seed virus was treated with 2 μl TURBO DNase (Thermo Fisher Scientific, Cat# AM2238) and incubated at 37°C for 1 h. Complete removal of the CPER products (i.e., SARS-CoV-2-related DNA) from the seed virus was verified by PCR. The working virus stock was prepared from the seed virus as described above.

To generate chimeric recombinant SARS-CoV-2 that encodes the S proteins of B.1.1, BA.1 and BA.2 (**Extended Data Fig. 4**), mutations were inserted in fragment 8 (**Supplementary Table 4**) using the GENEART site-directed mutagenesis system (Thermo Fisher Scientific, Cat# A13312) according to the manufacturer’s protocol with the primers listed in **Supplementary Table 5**. A recombinant SARS-CoV-2 that bears D614G S (corresponds to B.1.1 S) was prepared in our previous study^23^. To prepare a chimeric recombinant SARS-CoV-2 that bears Delta S (**Extended Data Fig. 4**), the fragment of viral genome that corresponds to the region of fragment 8 (**Supplementary Table 4**) was subcloned from a Delta isolate (strain TKYTK1734; GISAID ID: EPI_ISL_2378732)^23^. Nucleotide sequences were determined by a DNA sequencing service (Fasmac), and the sequence data were analysed by Sequencher software v5.1 (Gene Codes Corporation).

To produce chimeric recombinant SARS-CoV-2, the CPER products were transfected into HEK293-C34 cells using TransIT-LT1 (Takara, Cat# MIR2300) according to the manufacturer’s protocol. At 1 day posttransfection, the culture medium was replaced with Dulbecco’s modified Eagle’s medium (high glucose) (Sigma-Aldrich, Cat# R8758-500ML) containing 2% FCS, 1% PS and doxycycline (1 μg/ml; Takara, Cat# 1311N). At 7 days posttransfection, the culture medium was harvested and centrifuged, and the supernatants were collected as the seed virus. To remove the CPER products (i.e., SARS-CoV-2-related DNA), 1 ml of the seed virus was treated with 2 μl TURBO DNase (Thermo Fisher Scientific, Cat# AM2238) and incubated at 37°C for 1 h. Complete removal of the CPER products (i.e., SARS-CoV-2-related DNA) from the seed virus was verified by PCR. The working virus stock was prepared from the seed virus as described below (see “SARS-CoV-2 preparation and titration” section).

### SARS-CoV-2 preparation and titration

The working virus stocks of chimeric recombinant SARS-CoV-2 that encodes the S proteins of B.1.1, Delta, BA.1 and BA.2 were prepared and titrated as previously described^21–23^. In brief, 20 μl of the seed virus was inoculated into VeroE6/TMPRSS2 cells (5,000,000 cells in a T-75 flask). One hour after infection (h.p.i.), the culture medium was replaced with DMEM (low glucose) (Wako, Cat# 041-29775) containing 2% FBS and 1% PS. At 3 d.p.i., the culture medium was harvested and centrifuged, and the supernatants were collected as the working virus stock.

The titre of the prepared working virus was measured as the 50% tissue culture infectious dose (TCID_50_). Briefly, one day before infection, VeroE6/TMPRSS2 cells (10,000 cells) were seeded into a 96-well plate. Serially diluted virus stocks were inoculated into the cells and incubated at 37°C for 4 d. The cells were observed under microscopy to judge the CPE appearance. The value of TCID_50_/ml was calculated with the Reed–Muench method^45^.

To verify the sequence of chimeric recombinant SARS-CoV-2, viral RNA was extracted from the working viruses using a QIAamp viral RNA mini kit (Qiagen, Cat# 52906) and viral genome sequence was analysed as described above (see “Viral genome sequencing” section). In brief, the viral sequences of *GFP*-encoding recombinant SARS-CoV-2 (strain WK-521; GISIAD ID: EPI_ISL_408667)^21, 29^ that harbour the *S* genes of respective variants (B.1.1, BA.1 or BA.2) were used for the reference. Information on the unexpected mutations detected is summarized in **Supplementary Table 6**, and the raw data are deposited in Gene Expression Omnibus (accession number: GSE196649).

### SARS-CoV-2 infection

One day before infection, Vero cells (10,000 cells), VeroE6/TMPRSS2 cells (10,000 cells), Calu-3 cells (20,000 cells), HEK293-ACE2 cells (10,000 cells), HEK293-ACE2/TMPRSS2 cells (10,000 cells), were seeded into a 96-well plate. SARS-CoV-2 [100 TCID_50_ for VeroE6/TMPRSS2 cells (**Fig. 3a**), 1,000 TCID_50_ for Vero cells (**Fig. 3a**), HEK293-ACE2 cells (10,000 cells) (**Fig. 3j**), and HEK293-ACE2/TMPRSS2 cells (10,000 cells) (**Fig. 3j**); and 2,000 TCID_50_ for Calu-3 cells (**Fig. 3a**)] was inoculated and incubated at 37°C for 1 h. The infected cells were washed, and 180 µl of culture medium was added. The culture supernatant (10 µl) was harvested at the indicated timepoints and used for RT–qPCR to quantify the viral RNA copy number (see “RT–qPCR” section below).

The infection experiment primary human nasal epithelial cells (**Fig. 3a**) was performed as previously described^11, 23^. Briefly, the working viruses were diluted with Opti-MEM (Thermo Fisher Scientific, Cat# 11058021). The diluted viruses (1,000 TCID_50_ in 100 μl) were inoculated onto the apical side of the culture and incubated at 37°C for 1 h. The inoculated viruses were removed and washed twice with Opti-MEM. To harvest the viruses on the apical side of the culture, 100 μl Opti-MEM was applied onto the apical side of the culture and incubated at 37°C for 10 m. The Opti-MEM applied was harvested and used for RT–qPCR to quantify the viral RNA copy number (see “RT–qPCR” section below).

### RT–qPCR

RT–qPCR was performed as previously described^11, 12, 22, 23^. Briefly, 5 μl of culture supernatant was mixed with 5 μl of 2 × RNA lysis buffer [2% Triton X-100, 50 mM KCl, 100 mM Tris-HCl (pH 7.4), 40% glycerol, 0.8 U/μl recombinant RNase inhibitor (Takara, Cat# 2313B)] and incubated at room temperature for 10 min. RNase-free water (90 μl) was added, and the diluted sample (2.5 μl) was used as the template for real-time RT-PCR performed according to the manufacturer’s protocol using the One Step TB Green PrimeScript PLUS RT-PCR kit (Takara, Cat# RR096A) and the following primers: Forward *N*, 5’-AGC CTC TTC TCG TTC CTC ATC AC-3’; and Reverse *N*, 5’-CCG CCA TTG CCA GCC ATT C-3’. The viral RNA copy number was standardized with a SARS-CoV-2 direct detection RT-qPCR kit (Takara, Cat# RC300A). Fluorescent signals were acquired using QuantStudio 3 Real-Time PCR system (Thermo Fisher Scientific), CFX Connect Real-Time PCR Detection system (Bio-Rad), Eco Real-Time PCR System (Illumina), qTOWER3 G Real-Time System (Analytik Jena) or 7500 Real-Time PCR System (Thermo Fisher Scientific).

### Fluorescence microscopy

Fluorescence microscopy (**Fig. 3b** and **Extended Data Fig. 5a**) was performed as previously described^23^. Briefly, one day before infection, VeroE6/TMPRSS2 cells (10,000 cells) were seeded into 96-well, glass bottom, black plates and infected with SARS-CoV-2 (100 TCID_50_). At 24, 48, and 72 h.p.i., GFP fluorescence was observed under an All-in-One Fluorescence Microscope BZ-X800 (Keyence) in living cells, and a 13-square-millimeter area of each sample was scanned. Images were reconstructed using an BZ-X800 analyzer software (Keyence), and the area of the GFP-positive cells was measured using this software.

### Plaque assay

Plaque assay (**Fig. 3c** and **Extended Data Fig. 5b**) was performed as previously described^12, 22, 23^. Briefly, one day before infection, VeroE6/TMPRSS2 cells (100,000 cells) were seeded into a 24-well plate and infected with SARS-CoV-2 (1, 10, 100 and 1,000 TCID_50_) at 37°C for 2 h. Mounting solution containing 3% FBS and 1.5% carboxymethyl cellulose (Wako, Cat# 039-01335) was overlaid, followed by incubation at 37°C. At 3 d.p.i., the culture medium was removed, and the cells were washed with PBS three times and fixed with 4% paraformaldehyde phosphate (Nacalai Tesque, Cat# 09154-85). The fixed cells were washed with tap water, dried, and stained with staining solution [0.1% methylene blue (Nacalai Tesque, Cat# 22412-14) in water] for 30 m. The stained cells were washed with tap water and dried, and the size of plaques was measured using Fiji software v2.2.0 (ImageJ).

### Coculture experiment

Coculture experiment (**Extended Data Fig. 5c**) was performed as previously described^12^. Briefly, one day before transfection, effector cells (i.e., S-expressing cells) were seeded on the poly-L-lysine (Sigma, Cat# P4832) coated coverslips put in a 12-well plate, and target cells were prepared at a density of 100,000 cells in a 12-well plate. To prepare effector cells, HEK293 cells were cotransfected with the S-expression plasmids (500 ng) and pDSP_8-11_ (500 ng) using PEI Max (Polysciences, Cat# 24765-1). To prepare target cells, HEK293 and HEK293-ACE2/TMPRSS2 cells were transfected with pDSP_1-7_ (500 ng). At 24 h posttransfection, target cells were detached and cocultured with effector cells in a 1:2 ratio. At 9 h post-coculture, cells were fixed with 4% parafprmaldehyde in PBS (Nacalai Tesque, Cat# 09154-85) for 15 m at room temperature. Nuclei were stained with Hoechst 33342 (Thermo Fisher Scientific, Cat# H3570). The coverslips were mounted on glass slides using Fluoromount-G (Southern Biotechnology, Cat# 0100-01) with Hoechst 33342 and observed using an A1Rsi Confocal Microscope (Nikon). The size of syncytium (GFP-positive area) was measured using Fiji software v2.2.0 (ImageJ) as previously described^12^.

### SARS-CoV-2 S-based fusion assay

SARS-CoV-2 S-based fusion assay (**Fig. 3d and 3f**) was performed as previously described^12, 22, 23^. This assay utilizes a dual split protein (DSP) encoding *Renilla* luciferase and *GFP* genes; the respective split proteins, DSP_8-11_ and DSP_1-7_, are expressed in effector and target cells by transfection. Briefly, on day 1, effector cells (i.e., S-expressing cells) and target cells (see below) were prepared at a density of 0.6–0.8 × 10^6^ cells in a 6-well plate. To prepare effector cells, HEK293 cells were cotransfected with the S expression plasmids (400 ng) and pDSP_8-11_ (400 ng) using TransIT-LT1 (Takara, Cat# MIR2300). To prepare target cells, HEK293 cells were cotransfected with pC-ACE2 (200 ng) and pDSP_1-7_ (400 ng). Target HEK293 cells in selected wells were cotransfected with pC-TMPRSS2 (40 ng) in addition to the plasmids above. VeroE6/TMPRSS2 cells, Calu-3 cells, HEK293-ACE2 cells and HEK293-ACE2/TMPRSS2 cells were transfected with pDSP_1-7_ (400ng). On day 3 (24 h posttransfection), 16,000 effector cells were detached and reseeded into 96-well black plates (PerkinElmer, Cat# 6005225), and target cells (HEK293, VeroE6/TMPRSS2 or Calu-3/DSP_1-7_ cells) were reseeded at a density of 1,000,000 cells/2 ml/well in 6-well plates. On day 4 (48 h posttransfection), target cells were incubated with EnduRen live cell substrate (Promega, Cat# E6481) for 3 h and then detached, and 32,000 target cells were added to a 96-well plate with effector cells. *Renilla* luciferase activity was measured at the indicated time points using Centro XS3 LB960 (Berthhold Technologies). To measure the surface expression level of S protein, effector cells were stained with rabbit anti-SARS-CoV-2 S S1/S2 polyclonal antibody (Thermo Fisher Scientific, Cat# PA5-112048, 1:100). Normal rabbit IgG (SouthernBiotech, Cat# 0111-01, 1:100) was used as negative controls, and APC-conjugated goat anti-rabbit IgG polyclonal antibody (Jackson ImmunoResearch, Cat# 111-136-144, 1:50) was used as a secondary antibody. Surface expression level of S proteins (**Extended Data Fig. 6a**) was measured using FACS Canto II (BD Biosciences) and the data were analysed using FlowJo software v10.7.1 (BD Biosciences). Gating strategy for flow cytometry is shown in **Supplementary Fig. 1**. To calculate fusion activity, *Renilla* luciferase activity was normalized to the MFI of surface S proteins. The normalized value (i.e., *Renilla* luciferase activity per the surface S MFI) is shown as fusion activity.

### Yeast surface display

Yeast surface display (**Extended Data Fig. 6b**) was performed as previously described as previously described^22, 24, 46^. Briefly, the peptidase domain of human ACE2 (residues 18-740) was expressed in Expi293 cells and purified by a 5-ml HisTrap Fast Flow column (Cytiva, Cat# 17-5255-01) and Superdex 200 16/600 (Cytiva, Cat# 28-9893-35) using an ÄKTA pure chromatography system (Cytiva), and the purified soluble ACE2 was labelled with CF640 (Biotium, Cat# 92108). Protein quality was verified using a Tycho NT.6 system (NanoTemper) and ACE2 activity assay kit (SensoLyte, Cat# AS-72086).

An enhanced yeast display platform for SARS-CoV-2 RBD [wild-type (B.1.1), residues 336-528] yeast surface expression was established using *Saccharomyces cerevisiae* EBY100 strain and pJYDC1 plasmid (Addgene, Cat# 162458) as previously described^22, 24, 47^. The yeast-optimized SARS-CoV-2_RBD-Omicron-BA.1 gene (**Supplementary Table 7**) was obtained from Twist Biosciences. The site-directed mutagenesis was performed using the KAPA HiFi HotStart ReadyMix kit (Roche, Cat# KK2601) by restriction enzyme-free cloning procedure^48^. Primers for mutagenesis are listed in **Supplementary Table 8**.

The binding affinity of SARS-CoV-2 S B.1.1, BA.1, and BA.2 RBDs to human ACE2 were titrated by flow cytometry. The CF640-labelled ACE2 at 12–14 different concentrations (200 nM to 13 pM in PBS supplemented with bovine serum albumin at 1 g/l) per measurement were incubated with expressed yeast aliquots and 1 nM bilirubin (Sigma-Aldrich, Cat# 14370-1G) and analysed by using FACS S3e Cell Sorter device (Bio-Rad) as previously described^22, 24, 47^. The background binding subtracted fluorescent signal was fitted to a standard noncooperative Hill equation by nonlinear least-squares regression using Python v3.7 (https://www.python.org) as previously described^47^.

### TMPRSS2 expression on the cell surface

To measure the surface expression level of TMPRSS2 (**Extended Data Fig. 6c**), HEK293-ACE2 cells and HEK293-ACE2/TMPRSS2 cells were stained with rabbit anti-TMPRSS2 polyclonal antibody (BIOSS, Cat# BS-6285R, 1:100). Normal rabbit IgG (SouthernBiotech, Cat# 0111-01, 1:100) was used as negative controls, and APC-conjugated goat anti-rabbit IgG polyclonal antibody (Jackson ImmunoResearch, Cat# 111-136-144, 1:50) was used as a secondary antibody. Surface expression level of TMPRSS2 was measured using FACS Canto II (BD Biosciences) and the data were analysed using FlowJo software v10.7.1 (BD Biosciences). Gating strategy for flow cytometry is shown in **Supplementary Fig. 1**.

### Western blot

Western blot (**Fig. 3e**) was performed as previously described^12, 22, 23^. The HEK293 cells cotransfected with the S expression plasmids and pDSP_8-11_ (see “SARS-CoV-2 S-based fusion assay” section above) were used. To quantify the level of the cleaved S2 protein in the cells, the harvested cells were washed and lysed in lysis buffer [25 mM HEPES (pH 7.2), 20% glycerol, 125 mM NaCl, 1% Nonidet P40 substitute (Nacalai Tesque, Cat# 18558-54), protease inhibitor cocktail (Nacalai Tesque, Cat# 03969-21)]. After quantification of total protein by protein assay dye (Bio-Rad, Cat# 5000006), lysates were diluted with 2 × sample buffer [100 mM Tris-HCl (pH 6.8), 4% SDS, 12% β-mercaptoethanol, 20% glycerol, 0.05% bromophenol blue] and boiled for 10 m. Then, 10 μl samples (50 μg of total protein) were subjected to Western blot. For protein detection, the following antibodies were used: mouse anti-SARS-CoV-2 S monoclonal antibody (clone 1A9, GeneTex, Cat# GTX632604, 1:10,000), rabbit anti-beta actin (ACTB) monoclonal antibody (clone 13E5, Cell Signalling, Cat# 4970, 1:5,000), horseradish peroxidase (HRP)-conjugated donkey anti-rabbit IgG polyclonal antibody (Jackson ImmunoResearch, Cat# 711-035-152, 1:10,000) and HRP-conjugated donkey anti-mouse IgG polyclonal antibody (Jackson ImmunoResearch, Cat# 715-035-150, 1:10,000). Chemiluminescence was detected using SuperSignal West Femto Maximum Sensitivity Substrate (Thermo Fisher Scientific, Cat# 34095) according to the manufacturer’s instruction. Bands were visualized using an Amersham Imager 600 (GE Healthcare), and the band intensity was quantified using Image Studio Lite v5.2 (LI-COR Biosciences) or Fiji software v2.2.0 (ImageJ). Uncropped blots are shown in **Supplementary Fig. 2**.

### Pseudovirus infection

Pseudovirus infection was (**Fig. 3g**) performed as previously described^11, 12, 22–24, 43^. Briefly, the same amount of pseudoviruses (normalized to the HiBiT value, which indicates the amount of p24 HIV-1 antigen) was inoculated into HEK293-ACE2 cells or HEK293-ACE2/TMPRSS2 and viral infectivity was measured as described above (see “Neutralisation assay” section). To analyse the effect of TMPRSS2 for pseudovirus infectivity, the fold change of the values of HEK293-ACE2/TMPRSS2 to HEK293-ACE2 was calculated.

### Animal experiments

Animal experiments (**Fig. 4**) were performed as previously described^12, 23^. Syrian hamsters (male, 4 weeks old) were purchased from Japan SLC Inc. (Shizuoka, Japan). Baseline body weights were measured before infection. For the virus infection experiments, hamsters were euthanized by intramuscular injection of a mixture of either 0.15 mg/kg medetomidine hydrochloride (Domitor^®^, Nippon Zenyaku Kogyo), 2.0 mg/kg midazolam (Dormicum^®^, FUJIFILM Wako Chemicals) and 2.5 mg/kg butorphanol (Vetorphale^®^, Meiji Seika Pharma), or 0.15 mg/kg medetomidine hydrochloride, 2.0 mg/kg alphaxaone (Alfaxan^®^, Jurox) and 2.5 mg/kg butorphanol. The B.1.1 virus, Delta, Omicron (10,000 TCID_50_ in 100 µl), or saline (100 µl) were intranasally inoculated under anaesthesia. Oral swabs were daily collected under anaesthesia with isoflurane (Sumitomo Dainippon Pharma). Body weight, enhanced pause (Penh), the ratio of time to peak expiratory follow relative to the total expiratory time (Rpef) and subcutaneous oxygen saturation (SpO_2_) were routinely monitored at indicated timepoints (see “Lung function test” section below). Respiratory organs were anatomically collected at 1, 3 and 5 d.p.i (for lung) or 1 d.p.i. (for trachea). Viral RNA load in the respiratory tissues were determined by RT–qPCR. The respiratory tissues were also used for histopathological and IHC analyses (see “H&E staining” and “IHC” sections below).

### Lung function test

Lung function test (**Fig. 4a**) was performed as previously described^12^. Respiratory parameters (Penh and Rpef) were measured by using a whole-body plethysmography system (DSI) according to the manufacturer’s instructions. In brief, a hamster was placed in an unrestrained plethysmography chamber and allowed to acclimatize for 30 s, then, data were acquired over a 4-m period by using FinePointe Station and Review softwares v2.9.2.12849 (STARR). The state of oxygenation was examined by measuring SpO_2_ using pulse oximeter, MouseOx PLUS (STARR). SpO_2_ was measured by attaching a measuring chip to the neck of hamsters sedated by 0.25 mg/kg medetomidine hydrochloride.

### IHC

IHC (**Extended Data Fig. 7**) was performed as previously described^12, 23^ using an Autostainer Link 48 (Dako). The deparaffinized sections were exposed to EnVision FLEX target retrieval solution high pH (Agilent, Cat# K8004) for 20 m at 97°C to activate, and mouse anti-SARS-CoV-2 N monoclonal antibody (R&D systems, Clone 1035111, Cat# MAB10474-SP, 1:400) was used as a primary antibody. The sections were sensitized using EnVision FLEX (Agilent) for 15 m and visualised by peroxidase-based enzymatic reaction with 3,3’-diaminobenzidine tetrahydrochloride as substrate for 5 m. The N protein positivity (**Fig. 4c and 4d**) was evaluated by certificated pathologists as previously described^12^. Images were incorporated as virtual slide by NDRscan3.2 software (Hamamatsu Photonics). The N-protein positivity was measured as the area using Fiji software v2.2.0 (ImageJ).

### H&E staining

H&E staining (**Extended Data Fig. 8**) was performed as previously described^12, 23^. Briefly, excised animal tissues were fixed with 10% formalin neutral buffer solution, and processed for paraffin embedding. The paraffin blocks were sectioned with 3 µm-thickness and then mounted on silane-coated glass slides (MAS-GP, Matsunami). H&E staining was performed according to a standard protocol.

### Histopathological scoring

Histopathological scoring (**Fig. 4e** and **Extended Data Fig. 8a**) was performed as previously described^12, 23^. Pathological features including bronchitis or bronchiolitis, haemorrhage with congestive edema, alveolar damage with epithelial apoptosis and macrophage infiltration, hyperplasia of type II pneumocytes, and the area of the hyperplasia of large type II pneumocytes were evaluated by certified pathologists and the degree of these pathological findings were arbitrarily scored using four-tiered system as 0 (negative), 1 (weak), 2 (moderate), and 3 (severe). The “large type II pneumocytes” are the hyperplasia of type II pneumocytes exhibiting more than 10-μm-diameter nucleus. We described “large type II pneumocytes” as one of the remarkable histopathological features reacting SARS-CoV-2 infection in our previous studies^12, 23^. Total histology score is the sum of these five indices.

To measure the inflammation area in the infected lungs (**Extended Data Fig. 8b**), four hamsters infected with each virus were sacrificed at the 1, 3 and 5 d.p.i., and all four right lung lobes, including upper (anterior/cranial), middle, lower (posterior/caudal), and accessory lobes, were sectioned along with their bronchi. The tissue sections were stained by H&E, and the digital microscopic images were incorporated into virtual slides using NDRscan3.2 software (Hamamatsu Photonics). The inflammatory area including type II pneumocyte hyperplasia in the infected whole lungs was morphometrically analysed using Fiji software v2.2.0 (ImageJ).

### Statistics and reproducibility

Statistical significance was tested using a two-sided Student’s *t*-test or a two-sided Mann–Whitney *U*-test unless otherwise noted. The tests above were performed using Prism 9 software v9.1.1 (GraphPad Software).

In the time-course experiments (**Fig. 3a, 3d, 3f, 3h**, **4a–4c,** and **4e**), a multiple regression analysis including experimental conditions (i.e., the types of infected viruses) as explanatory variables and timepoints as qualitative control variables was performed to evaluate the difference between experimental conditions thorough all timepoints. *P* value was calculated by a two-sided Wald test. Subsequently, familywise error rates (FWERs) were calculated by the Holm method. These analyses were performed in v4.1.2 (https://www.r-project.org/).

In **Extended Data Fig. 7 and 8**, photographs shown are the representative areas of at least two independent experiments by using four hamsters at each timepoint. In **Extended Data Fig. 5**, assays were performed in triplicate. Photographs shown are the representatives of >20 fields of view taken for each sample.

## Data availability

The raw data of virus sequences analysed in this study are deposited in Gene Expression Omnibus (accession number: GSE196649). All databases/datasets used in this study are available from GISAID database (https://www.gisaid.org) and Genbank database (https://www.ncbi.nlm.nih.gov/genbank/). The accession numbers of viral sequences used in this study are listed in Method section.

## Code availability

The computational code to estimate the relative effective reproduction number of each viral lineage (**Fig. 1**) is available in the GitHub repository (https://github.com/TheSatoLab/Omicron_BA2/tree/main/lineage_growth_hierarchical_model).

## Author Contributions

Daichi Yamasoba, Izumi Kimura, Hesham Nasser, Keiya Uriu, Kotaro Shirakawa, Yusuke Kosugi, Mako Toyoda, Yuri L Tanaka, Erika P Butlertanaka, Ryo Shimizu, Kayoko Nagata, Takamasa Ueno, Akatsuki Saito, Takashi Irie, Terumasa Ikeda, Kei Sato performed cell culture experiments.

Jiri Zahradnik, Gideon Schreiber performed an yeast surface display assay.

Yuhei Morioka, Naganori Nao, Rigel Suzuki, Mai Kishimoto, Kouji Kobiyama, Teppei Hara, Hayato Ito, Yasuko Orba, Michihito Sasaki, Kumiko Yoshimatsu, Ken J Ishii, Hirofumi Sawa, Keita Matsuno, Takasuke Fukuhara performed animal experiments.

Masumi Tsuda, Lei Wang, Yoshitaka Oda, Shinya Tanaka performed histopathological analysis.

Hiroyuki Asakura, Mami Nagashima, Kenji Sadamasu, Kazuhisa Yoshimura performed viral genome sequencing analysis.

Kotaro Shirakawa, Jin Kuramochi, Motoaki Seki, Ryoji Fujiki, Atsushi Kaneda, Tadanaga Shimada, Taka-aki Nakada, Seiichiro Sakao, Takuji Suzuki, Akifumi Takaori-Kondo contributed clinical sample collection.

Jumpei Ito performed statistical, modelling, and bioinformatics analyses.

Jumpei Ito, Akatsuki Saito, Takashi Irie, Shinya Tanaka, Keita Matsuno, Takasuke Fukuhara, Terumasa Ikeda, and Kei Sato designed the experiments and interpreted the results.

Jumpei Ito and Kei Sato wrote the original manuscript.

All authors reviewed and proofread the manuscript.

The Genotype to Phenotype Japan (G2P-Japan) Consortium contributed to the project administration.

## Acknowledgments

We would like to thank all members belonging to The Genotype to Phenotype Japan (G2P-Japan) Consortium. We thank Dr. Kenzo Tokunaga (National Institute for Infectious Diseases, Japan) and Dr. Jin Gohda (The University of Tokyo, Japan) and Dr. Hisashi Arase (Osaka University) for providing reagents. The super-computing resource was provided by Human Genome Center at The University of Tokyo.

This study was supported in part by AMED Research Program on Emerging and Re-emerging Infectious Diseases (20fk0108268, to Akifumi Takaori-Kondo; 20fk0108517, to Akifumi Takaori-Kondo; 20fk0108401, to Takasuke Fukuhara; 20fk010847, to Takasuke Fukuhara; 21fk0108617 to Takasuke Fukuhara; 20fk0108146, to Kei Sato; 20fk0108270, to Kei Sato; and 20fk0108413, to Atsushi Kaneda, Terumasa Ikeda and Kei Sato) and (20fk0108451, to G2P-Japan Consortium, Akatsuki Saito, Takashi Irie, Keita Matsuno, Takasuke Fukuhara, Terumasa Ikeda, and Kei Sato); AMED Research Program on HIV/AIDS (21fk0410034, to Akifumi Takaori-Kondo; 21fk0410033, to Akatsuki Saito; and 21fk0410039, to Kei Sato); AMED CRDF Global Grant (21jk0210039 to Akatsuki Saito); AMED Japan Program for Infectious Diseases Research and Infrastructure (21wm0325009, to Akatsuki Saito; 21wm0125008, to Hirofumi Sawa and 21wm0225003, to Hirofumi Sawa); JST A-STEP (JPMJTM20SL, to Terumasa Ikeda); JST SICORP (e-ASIA) (JPMJSC20U1, to Kei Sato); JST SICORP (JPMJSC21U5, to Kei Sato), JST CREST (JPMJCR20H4, to Kei Sato); JSPS KAKENHI Grant-in-Aid for Scientific Research C (19K06382, to Akatsuki Saito); JSPS KAKENHI Grant-in-Aid for Scientific Research B (21H02736, to Takasuke Fukuhara; 18H02662, to Kei Sato; and 21H02737, to Kei Sato); JSPS Fund for the Promotion of Joint International Research (Fostering Joint International Research) (18KK0447, to Kei Sato); JSPS Core-to-Core Program (A. Advanced Research Networks) (JPJSCCA20190008, to Kei Sato); JSPS Research Fellow DC1 (19J20488, to Izumi Kimura); JSPS Leading Initiative for Excellent Young Researchers (LEADER) (to Terumasa Ikeda); World-leading Innovative and Smart Education (WISE) Program 1801 from the Ministry of Education, Culture, Sports, Science and Technology (MEXT) (to Naganori Nao); The Tokyo Biochemical Research Foundation (to Kei Sato); Mitsubishi Foundation (to Terumasa Ikeda); Shin-Nihon Foundation of Advanced Medical Research (to Mako Toyoda and Terumasa Ikeda); Tsuchiya Foundation (to Takashi Irie); a Grant for Joint Research Projects of the Research Institute for Microbial Diseases, Osaka University (to Akatsuki Saito); an intramural grant from Kumamoto University COVID-19 Research Projects (AMABIE) (to Terumasa Ikeda); Intercontinental Research and Educational Platform Aiming for Eradication of HIV/AIDS (to Terumasa Ikeda); and Joint Usage/Research Center program of Institute for Frontier Life and Medical Sciences, Kyoto University (to Kei Sato).

## Consortia

**The Genotype to Phenotype Japan (G2P-Japan) Consortium**

Mai Suganami^1^, Akiko Oide^1^, Mika Chiba^1^, Tomokazu Tamura^5^, Kana Tsushima^5^, Haruko Kubo^5^, Zannatul Ferdous^9^, Hiromi Mouri^9^, Miki Iida^9^, Keiko Kasahara^9^, Koshiro Tabata^9^, Mariko Ishizuka^9^, Asako Shigeno^29^, Kenzo Tokunaga^32^, Seiya Ozono^32^, Isao Yoshida^21^, So Nakagawa^33^, Jiaqi Wu^33^, Miyoko Takahashi^33^, Bahityar Rahmutulla Nawai^23^, Yutaka Suzuki^34^, Yukie Kashima^34^, Kazumi Abe^34^, Kiyomi Imamura^34^, Ryoko Kawabata^28^, Otowa Takahashi^3^, Kimiko Ichihara^3^, Kazuko Kitazato^3^, Haruyo Hasebe^3^, Chihiro Motozono^17^, Toong Seng Tan^17^, Isaac Ngare^17^

^32^ National Institute of Infectious Diseases, Tokyo, Japan

^33^ Tokai University, Isehara, Japan

^34^ The University of Tokyo, Kashiwa, Japan

---

**Extended Data Fig. 1.**
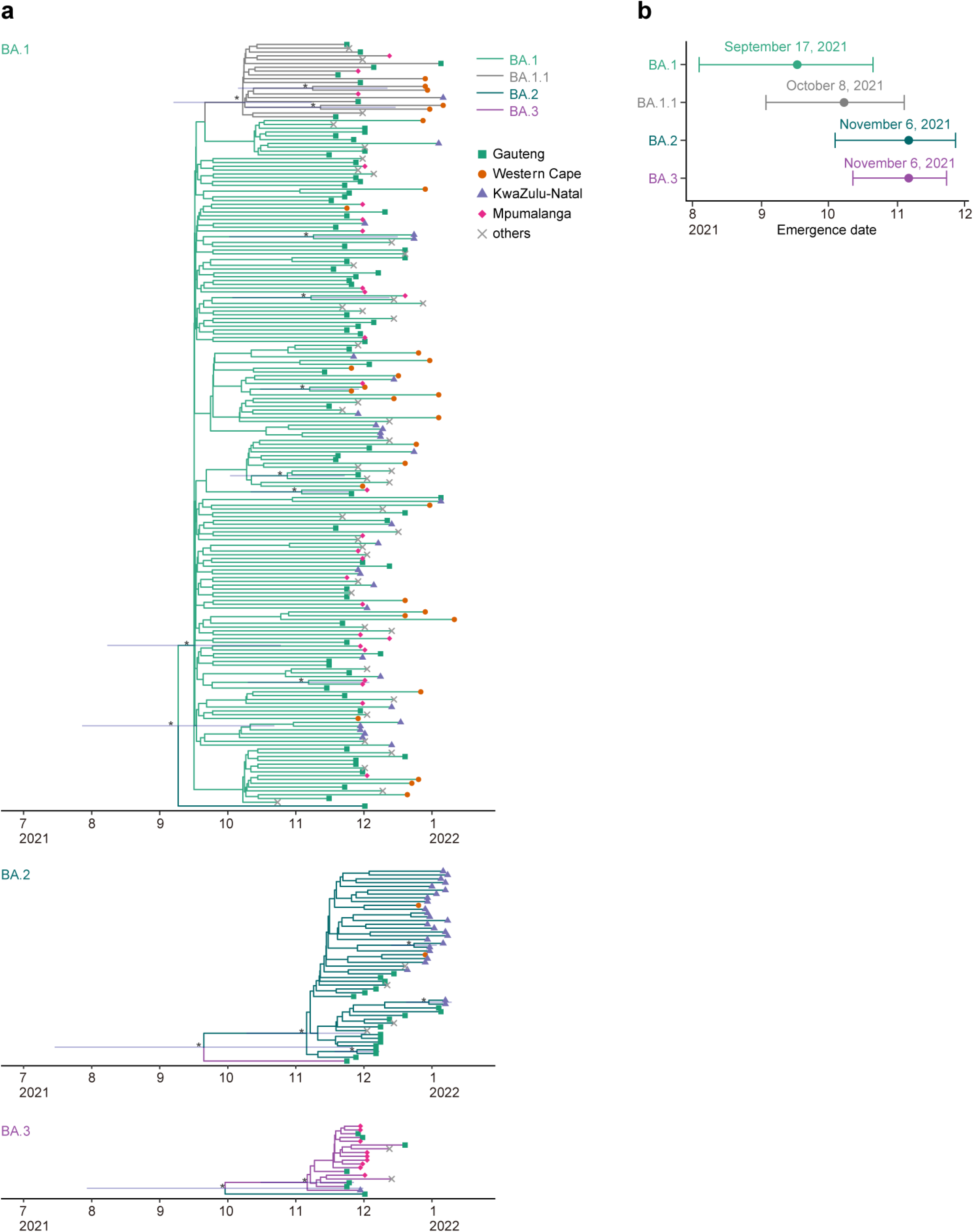
Estimated emergence dates of the Omicron lineages. **a**, Phylodynamics of BA.1 (top), BA.2 (middle), and BA.3 (bottom) sampled by January 26, 2022 in South Africa. All the BA.2 and BA.3 sequences and 200 randomly sampled BA.1 (including 20 BA.1.1) sequences were used. The time-resolved trees were constructed by BEAST2. Regarding a node with ≥0.95 posterior value (denoted by an asterisk), the 95% CI of the divergence time is shown. **b**, Estimated emergence dates of the Omicron lineages. The 95% CI (error bar) and posterior mean (dot) are shown.

**Extended Data Fig. 2.**
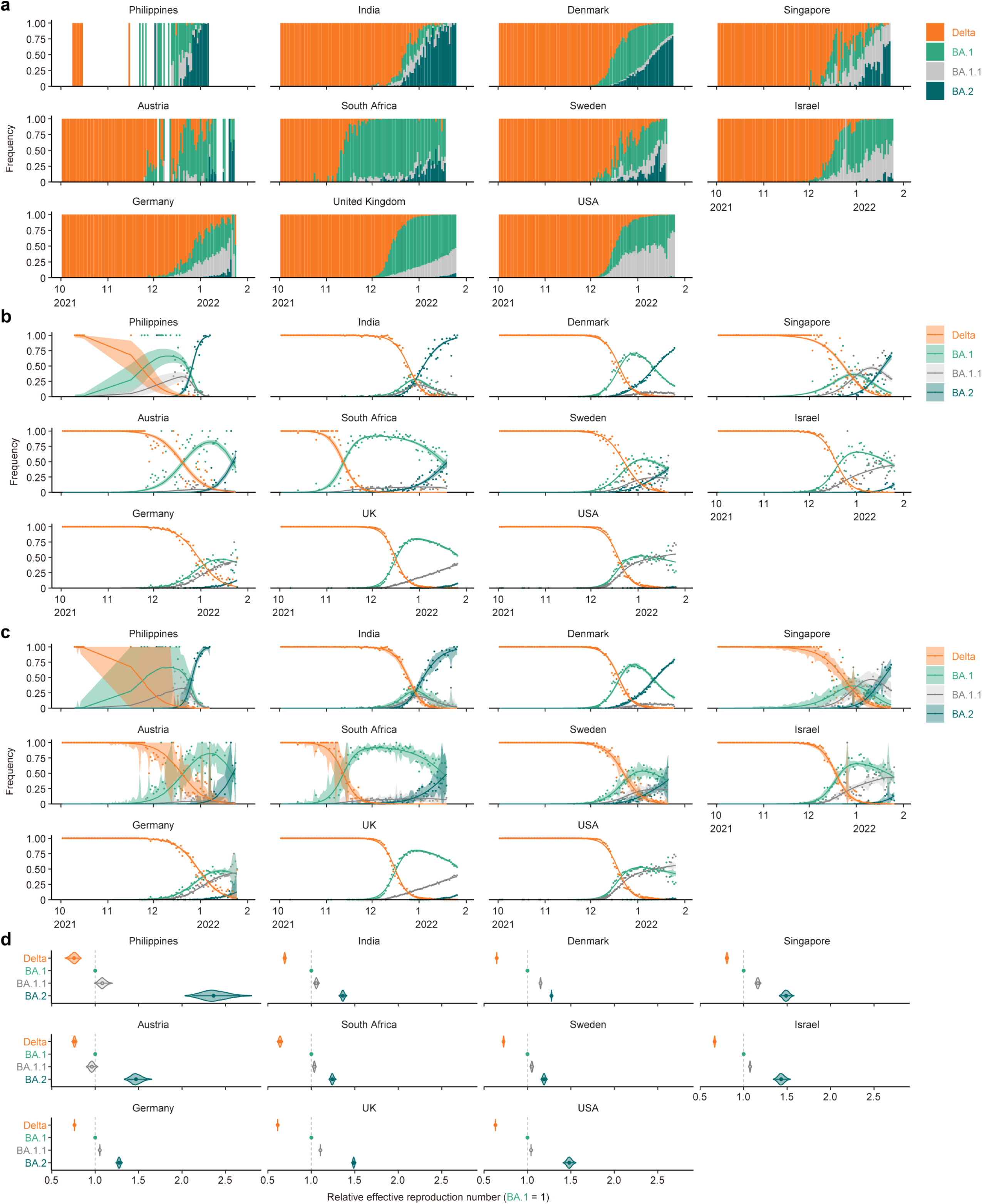
Epidemic dynamics of SARS-CoV-2 lineages in countries with the BA.2 epidemic. **a,** Daily sequence frequency of each viral lineage in eleven countries where ≥100 BA.2 sequences have been reported by January 25, 2022. This data was used as an input of a Bayesian hierarchical model to estimate the epidemic dynamics. **b,c,** Epidemic dynamics of SARS-CoV-2 viral lineages. The observed daily frequency (dot) and the dynamics estimated by the Bayesian model (posterior mean; line) are shown. Additionally, 95% CI (**b**) and 90% prediction interval (**c**) are shown. **d,** Estimated relative effective reproduction number of each viral lineage in each country. The posterior distribution (violin), 95% CI (line), and posterior mean (dot) are shown.

**Extended Data Fig. 3.**
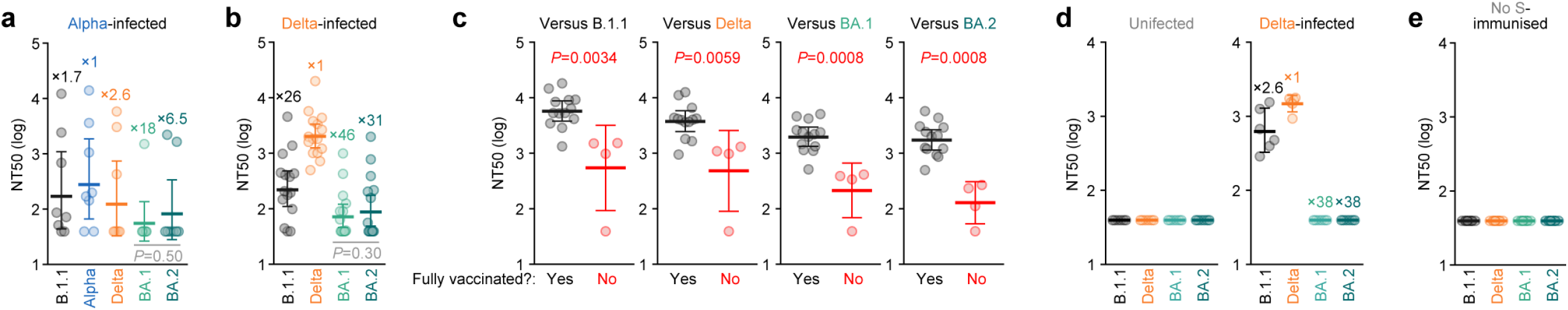
Immune resistance of BA.2. Neutralisation assays were performed with the pseudoviruses harbouring the S proteins of B.1.1 (the D614G-bearing ancestral virus), Alpha, Delta, BA.1 and BA.2. Assay of each serum sample was performed in triplicate to determine NT50, and each dot represents each NT50 value. Geometric mean and 95% CI are shown. The number indicates the fold change of resistance versus each antigenic variant. In **a** and **b**, statistically significant differences between BA.1 and BA.2 were determined by two-sided Wilcoxon signed-rank test. In **c**, statistically significant differences between fully-vaccinated (13 donors) and not-fully-vaccinated (4 donors) were determined by two-sided Mann-Whitney U-test. Information of vaccinated and convalescent donors are summarized in **Supplementary Tables 1 and 2**, respectively.

**Extended Data Fig. 4.**
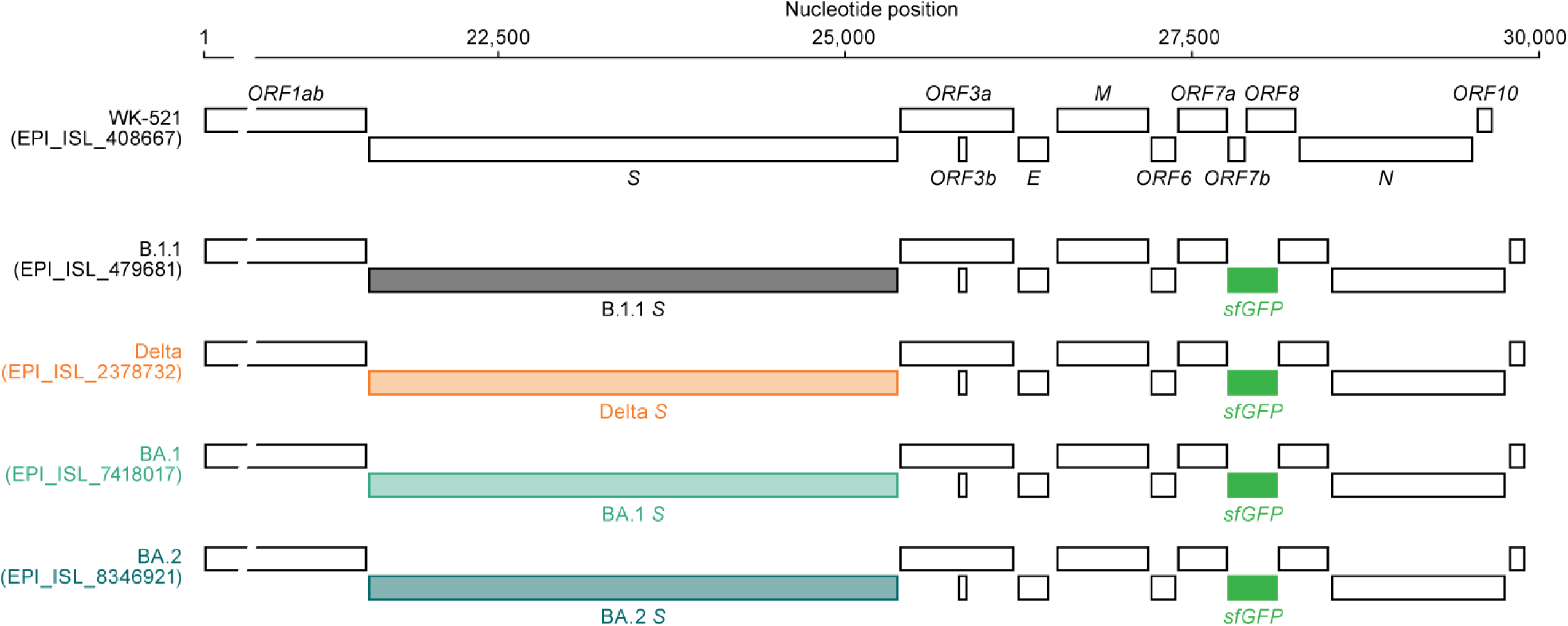
Scheme of chimeric recombinant SARS-CoV-2 used in this study. The SARS-CoV-2 genome and respective genes are shown. The template is SARS-CoV-2 strain WK-521 (lineage A, GISAID ID: EPI_ISL_408667), and the *S* genes were swapped with respective lineage/strain (GISAID IDs are indicated in the figure). Additionally, the *ORF7b* was swapped with *sfGFP* gene.

**Extended Data Fig. 5.**
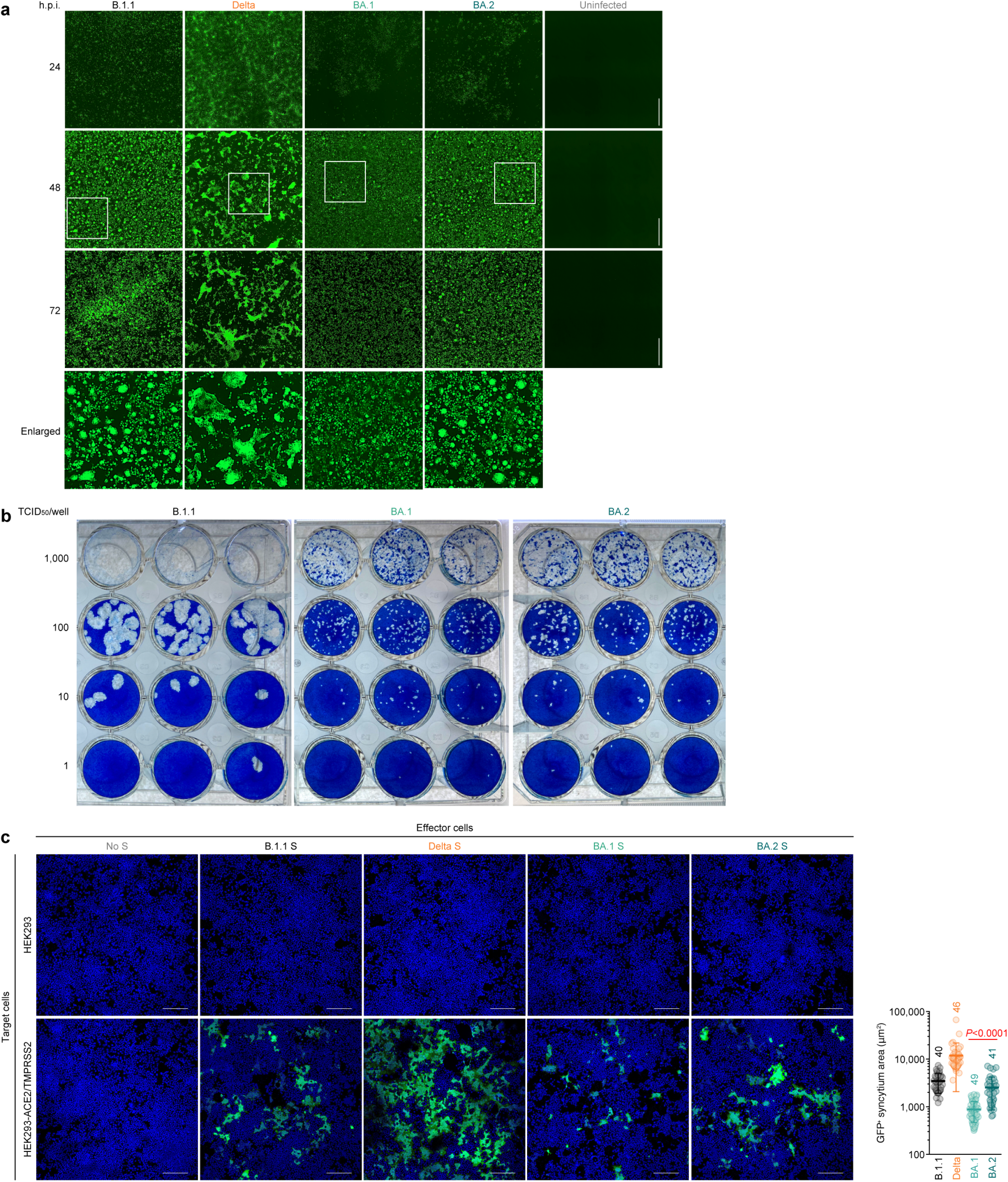
Syncytia and plaque formations by BA.2. **a,** Fluorescence microscopy. GFP area of infected VeroE6/TMPRSS2 cells (m.o.i. 0.01) at 24, 48, and 72 h.p.i were measured. Higher-magnification views of the regions indicated by squares are shown at bottom. **b**, Plaque assay. **c**, Coculture of S-expressing cells with HEK293-ACE2/TMPRSS2 cells. Left, representative images of S-expressing cells cocultured with HEK293 cells (top) or HEK293-ACE2/TMPRSS2 cells (bottom). Nuclei were stained with Hoechst 33342 (blue). Right, the size distribution of syncytia (green). Numbers in the panel indicate the numbers of GFP-positive syncytia counted. Data are the average ± s.d. A statistically significant difference between BA.1 an BA.2 was determined by two-sided Mann–Whitney *U*-test. In **a** and **b**, summarized data are shown in Fig. 3b and 3c. Scale bars, 500 μm (**a**) or 200 μm (**c**).

**Extended Data Fig. 6.**
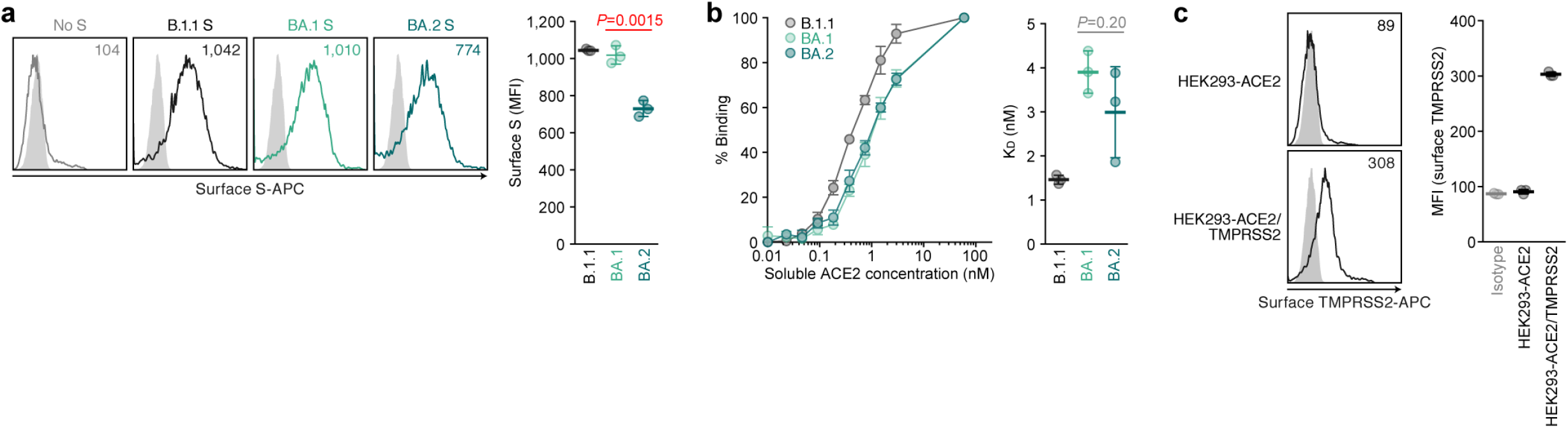
Expression and binding properties of BA.2 S. **a**, S expression on the cell surface. Representative histograms stained with anti-S1/S2 polyclonal antibody (left) and the summarised data (right) are respectively shown. The number in the histogram indicates MFI. Grey histograms indicate isotype controls. **b**, Binding affinity of SARS-CoV-2 S RBD to ACE2 by yeast surface display. Left, The percentage of the binding of the SARS-CoV-2 S RBD expressed on yeast to soluble ACE2 (left) and the summarised data (right) are respectively shown. **c**, TMPRSS2 expression on the cell surface. Left, representative histograms stained with anti-TMPRSS2 polyclonal antibody are shown. The number in the histogram indicates MFI. Grey histograms indicate isotype controls. Right, summarized data. Assays were performed in triplicate, and data are the average ± s.d. Each dot indicates the result from an individual replicate. A statistically significant difference between BA.1 an BA.2 was determined by two-sided unpaired Student’s *t*-test (**a**) or two-sided Mann–Whitney *U*-test (**b**).

**Extended Data Fig. 7.**
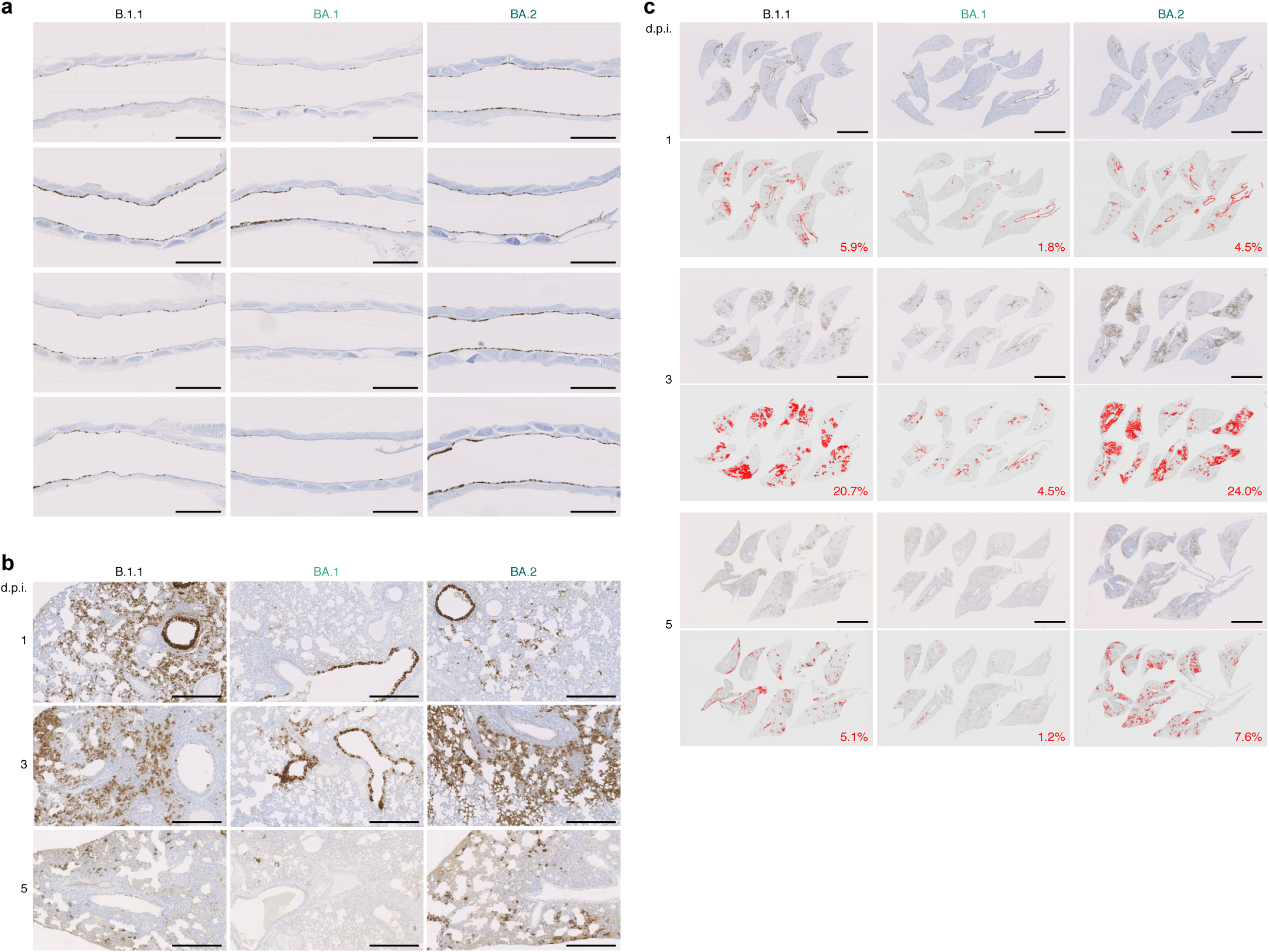
IHC of SARS-CoV-2 N protein. **a,b,** IHC of viral N protein in the middle part of trachea of all infected hamsters (n = 4 per each viral strain) at 1 d.p.i. (**a**) and the lung at 1, 3 and 5 d.p.i (**b**). Each panel indicates the representative result from an individual infected hamster. **c**, Lung lobes of the hamsters infected with B.1.1, BA.1 or BA.2 (n = 4 for each virus) at 1, 3 and 5 d.p.i. were immunohistochemically stained with anti-SARS-CoV-2 N monoclonal antibody. In each panel, IHC staining (top) and the digitalized N-positive area (bottom, indicated in red) are shown. The number in the bottom panel indicates the percentage of N-positive area. Summarized data is shown in Fig. 4c. Scale bars, 1 mm (**a**); 500 μm (**b**); or 5 mm (**c**).

**Extended Data Fig. 8.**
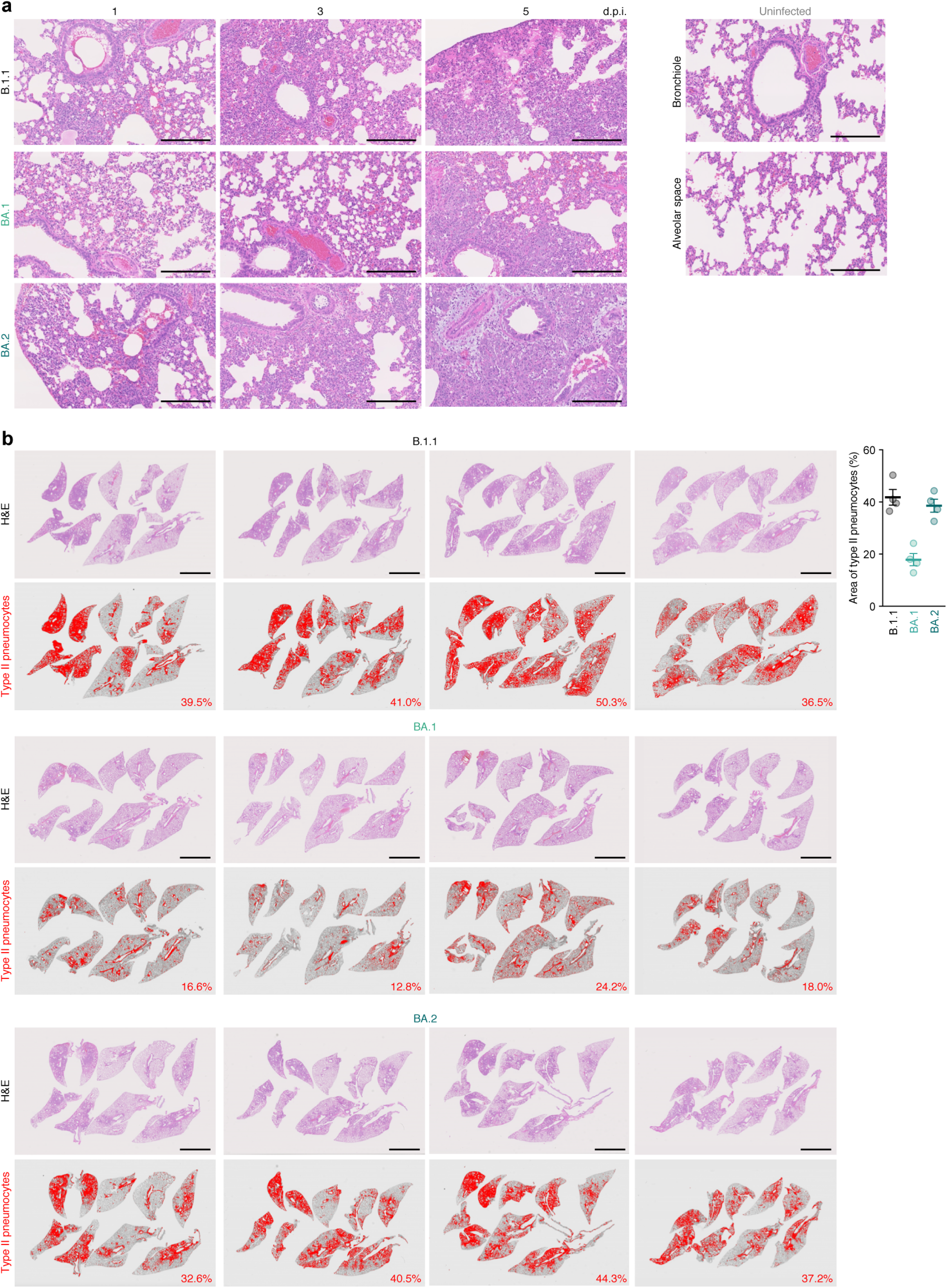
Pathological features of BA.2. **a**, H&E staining of the lungs of infected hamsters. Uninfected lung alveolar space and bronchioles are also shown. **b**, Type II pneumocytes in the lungs of infected hamsters. Left, lung lobes of the hamsters infected with B.1.1 (n = 4), BA.1 (n = 4), and BA.2 (n = 4) at 5 d.p.i. In each panel, H&E staining (top) and the digitalized inflammation area (bottom, indicated in red) are shown. The number in the bottom panel indicates the percentage of the section represented by the indicated area (i.e., the area indicated with red colour per the total area of the lung lobes). Right, summarized data. Each dot indicates the result from an individual hamster. Scale bars, 250 μm (**a**) or 5 mm (**b**).

## References

1. WHO. “Classification of Omicron (B.1.1.529): SARS-CoV-2 variant of concern (November 26, 2021)” https://www.who.int/news/item/26-11-2021-classification-of-omicron-(b.1.1.529)-sars-cov-2-variant-of-concern. (2021).

2. WHO. “Tracking SARS-CoV-2 variants (February 3, 2022)”. https://www.who.int/en/activities/tracking-SARS-CoV-2-variants/. (2022).

3. UKHSA. “SARS-CoV-2 variants of concern and variants under investigation in England. Technical briefing 35 (January 28, 2022)”. https://assets.publishing.service.gov.uk/government/uploads/system/uploads/attachment_data/file/1050999/Technical-Briefing-35-28January2022.pdf. (2022).

4. Cele, S. et al. Omicron extensively but incompletely escapes Pfizer BNT162b2 neutralization. Nature, doi: https://doi.org/10.1038/d41586-41021-03824-41585 (2021).

5. Cao, Y. et al. Omicron escapes the majority of existing SARS-CoV-2 neutralizing antibodies. Nature, doi: https://doi.org/10.1038/d41586-41021-03796-41586 (2021).

6. Dejnirattisai, W. et al. Reduced neutralisation of SARS-CoV-2 omicron B.1.1.529 variant by post-immunisation serum. Lancet, doi:https://doi.org/10.1016/S0140-6736(1021)02844-02840 (2021).

7. Cameroni, E. et al. Broadly neutralizing antibodies overcome SARS-CoV-2 Omicron antigenic shift. Nature, doi: https://doi.org/10.1038/d41586-41021-03825-41584 (2021).

8. Garcia-Beltran, W. F. et al. mRNA-based COVID-19 vaccine boosters induce neutralizing immunity against SARS-CoV-2 Omicron variant. Cell, doi: https://doi.org/10.1016/j.cell.2021.1012.1033 (2021).

9. Planas, D. et al. Considerable escape of SARS-CoV-2 Omicron to antibody neutralization. Nature, doi: https://doi.org/10.1038/d41586-41021-03827-41582 (2021).

10. Liu, L. et al. Striking antibody evasion manifested by the Omicron variant of SARS-CoV-2. Nature, doi: https://doi.org/10.1038/d41586-41021-03826-41583 (2021).

11. Meng, B. et al. Altered TMPRSS2 usage by SARS-CoV-2 Omicron impacts tropism and fusogenicity. Nature, doi:10.1038/s41586-022-04474-x (2022).

12. Suzuki, R. et al. Attenuated fusogenicity and pathogenicity of SARS-CoV-2 Omicron variant. Nature, doi:10.1038/s41586-022-04462-1 (2022).

13. Shuai, H. et al. Attenuated replication and pathogenicity of SARS-CoV-2 B.1.1.529 Omicron. Nature, doi:10.1038/s41586-022-04442-5 (2022).

14. Halfmann, P. J. et al. SARS-CoV-2 Omicron virus causes attenuated disease in mice and hamsters. Nature, doi:10.1038/s41586-022-04441-6 (2022).

15. Han, P. et al. Receptor binding and complex structures of human ACE2 to spike RBD from omicron and delta SARS-CoV-2. Cell, doi:10.1016/j.cell.2022.01.001 (2022).

16. Dejnirattisai, W. et al. SARS-CoV-2 Omicron-B.1.1.529 leads to widespread escape from neutralizing antibody responses. Cell 185, 467–484 e415, doi:10.1016/j.cell.2021.12.046 (2022).

17. Cameroni, E. et al. Broadly neutralizing antibodies overcome SARS-CoV-2 Omicron antigenic shift. Nature, doi:10.1038/s41586-021-04386-2 (2021).

18. Takashita, E. et al. Efficacy of Antibodies and Antiviral Drugs against Covid-19 Omicron Variant. N Engl J Med, doi:10.1056/NEJMc2119407 (2022).

19. VanBlargan, L. A. et al. An infectious SARS-CoV-2 B.1.1.529 Omicron virus escapes neutralization by therapeutic monoclonal antibodies. Nat Med, doi:10.1038/s41591-021-01678-y (2022).

20. Viana, R. et al. Rapid epidemic expansion of the SARS-CoV-2 Omicron variant in southern Africa. Nature, doi:10.1038/s41586-022-04411-y (2022).

21. Torii, S. et al. Establishment of a reverse genetics system for SARS-CoV-2 using circular polymerase extension reaction. Cell Rep 35, 109014 (2021).

22. Motozono, C. et al. SARS-CoV-2 spike L452R variant evades cellular immunity and increases infectivity. Cell Host Microbe 29, 1124–1136, doi:10.1016/j.chom.2021.06.006 (2021).

23. Saito, A. et al. Enhanced fusogenicity and pathogenicity of SARS-CoV-2 Delta P681R mutation. Nature, doi:10.1038/s41586-021-04266-9 (2021).

24. Kimura, I. et al. The SARS-CoV-2 Lambda variant exhibits enhanced infectivity and immune resistance. Cell Rep, doi: https://doi.org/10.1016/j.celrep.2021.110218 (2021).

25. Schubert, M. et al. Human serum from SARS-CoV-2 vaccinated and COVID-19 patients shows reduced binding to the RBD of SARS-CoV-2 Omicron variant. MedRxiv, doi: https://doi.org/10.1101/2021.1112.1110.21267523 (2022).

26. Wu, L. et al. SARS-CoV-2 Omicron RBD shows weaker binding affinity than the currently dominant Delta variant to human ACE2. Signal Transduct Target Ther 7, 8, doi:10.1038/s41392-021-00863-2 (2022).

27. Mlcochova, P. et al. SARS-CoV-2 B.1.617.2 Delta variant replication and immune evasion. Nature 599, 114–119, doi:10.1038/s41586-021-03944-y (2021).

28. Chen, S., Zhou, Y., Chen, Y. & Gu, J. fastp: an ultra-fast all-in-one FASTQ preprocessor. Bioinformatics 34, i884–i890, doi:10.1093/bioinformatics/bty560 (2018).

29. Matsuyama, S. et al. Enhanced isolation of SARS-CoV-2 by TMPRSS2-expressing cells. Proc Natl Acad Sci U S A 117, 7001–7003, doi:10.1073/pnas.2002589117 (2020).

30. Li, H. & Durbin, R. Fast and accurate short read alignment with Burrows-Wheeler transform. Bioinformatics 25, 1754–1760, doi:10.1093/bioinformatics/btp324 (2009).

31. Li, H. et al. The Sequence Alignment/Map format and SAMtools. Bioinformatics 25, 2078–2079, doi:10.1093/bioinformatics/btp352 (2009).

32. Cingolani, P. et al. A program for annotating and predicting the effects of single nucleotide polymorphisms, SnpEff: SNPs in the genome of Drosophila melanogaster strain w1118; iso-2; iso-3. Fly (Austin) 6, 80–92, doi:10.4161/fly.19695 (2012).

33. Li, H. Minimap2: pairwise alignment for nucleotide sequences. Bioinformatics 34, 3094–3100, doi:10.1093/bioinformatics/bty191 (2018).

34. Capella-Gutierrez, S., Silla-Martinez, J. M. & Gabaldon, T. trimAl: a tool for automated alignment trimming in large-scale phylogenetic analyses. Bioinformatics 25, 1972–1973, doi:10.1093/bioinformatics/btp348 (2009).

35. Stamatakis, A. RAxML version 8: a tool for phylogenetic analysis and post-analysis of large phylogenies. Bioinformatics 30, 1312–1313, doi:10.1093/bioinformatics/btu033 (2014).

36. Bouckaert, R. et al. BEAST 2: a software platform for Bayesian evolutionary analysis. PLoS Comput Biol 10, e1003537, doi:10.1371/journal.pcbi.1003537 (2014).

37. Hasegawa, M., Kishino, H. & Yano, T. Dating of the human-ape splitting by a molecular clock of mitochondrial DNA. J Mol Evol 22, 160–174, doi:10.1007/BF02101694 (1985).

38. Vohringer, H. S. et al. Genomic reconstruction of the SARS-CoV-2 epidemic in England. Nature 600, 506–511, doi:10.1038/s41586-021-04069-y (2021).

39. Obermeyer, F. et al. Analysis of 2.1 million SARS-CoV-2 genomes identifies mutations associated with transmissibility. MedRxiv, doi: https://doi.org/10.1101/2021.1109.1107.21263228 (2022).

40. Yamamoto, M. et al. The Anticoagulant Nafamostat Potently Inhibits SARS-CoV-2 S Protein-Mediated Fusion in a Cell Fusion Assay System and Viral Infection In Vitro in a Cell-Type-Dependent Manner. Viruses 12, doi:10.3390/v12060629 (2020).

41. Ozono, S. et al. SARS-CoV-2 D614G spike mutation increases entry efficiency with enhanced ACE2-binding affinity. Nat Commun 12, 848, doi:10.1038/s41467-021-21118-2 (2021).

42. Liu, Y. et al. The SARS-CoV-2 Delta variant is poised to acquire complete resistance to wild-type spike vaccines. BioRxiv, doi: https://doi.org/10.1101/2021.1108.1122.457114 (2021).

43. Uriu, K. et al. Neutralization of the SARS-CoV-2 Mu Variant by Convalescent and Vaccine Serum. N Engl J Med 385, 2397–2399, doi:10.1056/NEJMc2114706 (2021).

44. Ozono, S., Zhang, Y., Tobiume, M., Kishigami, S. & Tokunaga, K. Super-rapid quantitation of the production of HIV-1 harboring a luminescent peptide tag. J Biol Chem 295, 13023–13030, doi:10.1074/jbc.RA120.013887 (2020).

45. Reed, L. J. & Muench, H. A Simple Method of Estimating Fifty Percent Endpoints. Am J Hygiene 27, 493–497 (1938).

46. Zahradnik, J. et al. A Protein-Engineered, Enhanced Yeast Display Platform for Rapid Evolution of Challenging Targets. ACS Synth Biol 10, 3445–3460, doi:10.1021/acssynbio.1c00395 (2021).

47. Zahradnik, J. et al. SARS-CoV-2 variant prediction and antiviral drug design are enabled by RBD in vitro evolution. Nat Microbiol 6, 1188–1198, doi:10.1038/s41564-021-00954-4 (2021).

48. Peleg, Y. & Unger, T. Application of the restriction-free (RF) cloning for multicomponents assembly. Methods Mol Biol 1116, 73–87, doi:10.1007/978-1-62703-764-8_6 (2014).

