## Supplementary Figures for "Virological characteristics of SARS-CoV-2 BA.2 variant"

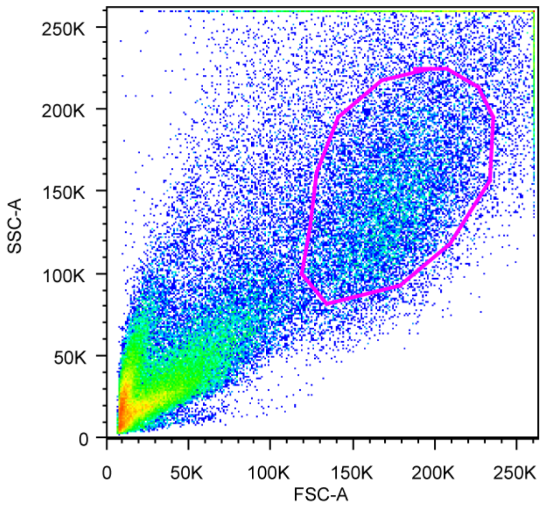


**Supplementary Fig. 1. Gating strategy for flow cytometry.**
A representative gating strategy for flow cytometry (**Extended Data Fig. 6b, c**) is shown.

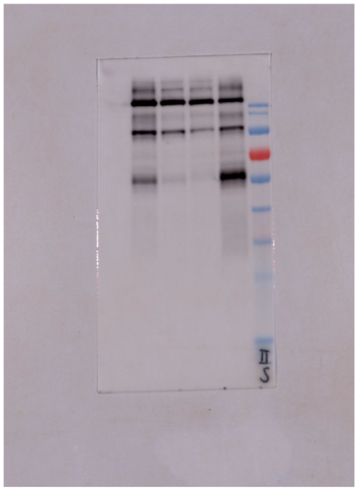

Mouse anti-SARS-CoV-2 S
monoclonal antibody
(clone 1A9, GeneTex,
Cat# GTX632604,
1:10,000)

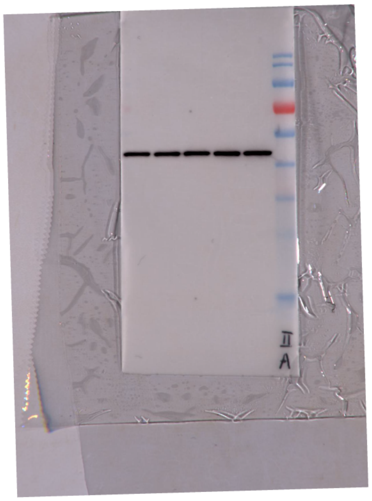

Rabbit anti-beta actin (ACTB)
monoclonal antibody
(clone 13E5, Cell Signalling,
Cat# 4970, 1:5,000)

**Supplementary Fig. 2. Original (uncrossed) blots.**
Uncrossed blots of **Fig. 3e** are shown.
